# Longitudinal associations between language network characteristics in the infant brain and school-age reading abilities are mediated by early-developing phonological skills

**DOI:** 10.1101/2023.06.22.546194

**Authors:** Xinyi Tang, Ted K. Turesky, Elizabeth S. Escalante, Megan Yf Loh, Mingrui Xia, Xi Yu, Nadine Gaab

**Author notes:** Corresponding author: Xi Yu, PhD, State Key Laboratory of Cognitive Neuroscience and Learning Beijing Normal University, Beijing 100875, China, 86-10-58805066.

## Abstract

Reading acquisition is a prolonged learning process relying on language development starting in utero. Behavioral longitudinal studies reveal prospective associations between infant language abilities and preschool/kindergarten phonological development that relates to subsequent reading performance. While recent pediatric neuroimaging work has begun to characterize the neural network underlying language development in infants, how this neural network scaffolds long-term language and reading acquisition remains unknown. We addressed this question in a 7-year longitudinal study from infancy to school-age. Seventy-six infants completed resting-state fMRI scanning, and underwent standardized language assessments in kindergarten. Of this larger cohort, forty-one were further assessed on their emergent word reading abilities after receiving formal reading instructions. Hierarchical clustering analyses identified a modular infant language network in which functional connectivity (FC) of the inferior frontal module prospectively correlated with kindergarten-age phonological skills and emergent word reading abilities. These correlations were obtained when controlling for infant age at scan, nonverbal IQ and parental education. Furthermore, kindergarten-age phonological skills mediated the relationship between infant FC and school-age reading abilities, implying a critical mid-way milestone for long-term reading development from infancy. Overall, our findings illuminate the neurobiological mechanisms by which infant language capacities could scaffold long-term reading acquisition.

**Highlights:**

- Clustering analyses revealed a modular language network in the infant brain
- Infant language network characteristics associate with school-age reading outcomes
- These longitudinal associations are mediated by kindergarten-age phonological skills

## 1. Introduction

Learning to read relies on language development starting in utero. Newborns show speech perception abilities (Gervain and Mehler, 2010), such as recognizing speech over non-speech sounds with comparable spectral and temporal characteristics (Vouloumanos and Werker, 2007) and discriminating phonetic differences in any language (e.g., descending versus ascending pitch contour, Nazzi et al., 1998). During the second half of the first year, infants gradually develop reliable phonemic representations of their native languages. They become adept at categorizing native-language phonetic units despite acoustic variations in specific phoneme exemplars produced by different speakers, while their abilities to differentiate nonnative phonemes decline, a phenomenon known as perceptual narrowing (Kuhl et al., 2006; Werker and Tees, 1984). Early linguistic experiences are theorized to serve as building blocks for various language skills, including phonological processing, in which children perceive and manipulate sound structures of words (Metsala and Walley, 2013; Vihman, 2014). Oral language skills that typically develop before schooling, in turn, are further postulated to lay critical foundations for the acquisition of word reading abilities. Thanks to the behavioral longitudinal investigations, prospective associations between preschool/kindergarten oral language skills, such as vocabulary size and phonological processing skills, and school-age reading performance have been repeatedly demonstrated to date (Hjetland et al., 2020; Hulme et al., 2015; Puolakanaho et al., 2008). By contrast, only a limited number of studies have explored the early trajectories of reading development in infancy and toddlerhood due to methodological challenges in infant behavioral assessments and longitudinal data acquisition. In these studies, infants’ language abilities, measured by expressive vocabulary size using the Early Language Test (Silvén, 1996) or MacArthur Communicative Development Inventories (CDI; Fenson et al., 1994), were shown to be associated with phonological skills in preschool; however, their specific contributions to school-age reading skills remain inconclusive (Silvén et al., 2004, 2007; Torppa et al., 2010). Empirical investigations are thus warranted to further unveil the mechanisms by which language development in infancy scaffolds long-term (pre-)reading acquisition.

Pediatric magnetic resonance imaging (MRI) techniques permit large-scale functional brain mappings in infants and children (Azhari et al., 2020; Graham et al., 2015). They enable the characterizing of the specific brain regions and their neural characteristics involved in language and reading processing across various developmental stages, providing valuable insights into the neurobiological mechanisms of the developmental trajectory of reading abilities. Our previous work (Yu et al., 2018) has shown that learning to read in school-age children is associated with increasingly stronger functional connectivity (FC) among the left occipitotemporal visual pathways and the bilateral frontotemporal regions that support language processing in both developing (Weiss-Croft and Baldeweg, 2015) and adult populations (Fedorenko et al., 2010). Moreover, while greater responses for written words compared to other visual categories, such as tools and faces, are only observed in the left-hemispheric visual pathway after reading onset (Dehaene-Lambertz et al., 2018), emerging evidence from task-based infant neuroimaging studies has revealed that infants exhibit adult-like functional characteristics in regions constituting the language network, such as bilateral inferior frontal and middle temporal regions (Dehaene-Lambertz, 2017). For example, infants have shown similar temporal organization of bilateral cortical response during sentence perception to that of adults (Dehaene-Lambertz et al., 2006), and both preverbal infants and adults have exhibited speech-sensitive activations in the bilateral inferior frontal and lateral temporal brain areas with stronger responses in the left hemisphere (May et al., 2018; Shultz et al., 2014; Vouloumanos et al., 2001). These bilateral regions further display intrinsic interhemispheric and intrahemispheric functional connections in sleeping infants (Smyser et al., 2016), a pattern that is also observable in the language network of adults (Braga et al., 2020; Tomasi and Volkow, 2012). Moreover, the emergent language neural network in infancy appeared to contain a frontal module and a temporal module, both of which showed greater FC for within-module connections than between-module connections, suggesting potentially differential developmental characteristics for each module (King et al., 2021). Longitudinally, FC patterns of key components of the bilateral language network in infancy are associated with children’s expressive and receptive language abilities in preschool (Emerson et al., 2016), as well as their phonological processing skills in kindergarten (Yu et al., 2021), highlighting the cognitive relevance of the functional organization of the infant brain. However, it is still unknown whether and how early-emerging language network characteristics in infancy *per se* relate to individual differences in children’s reading achievements at school age.

Utilizing a seven-year longitudinal dataset spanning infancy to school-ages, the current study investigated the long-term associations between FC in the emergent language network in infancy and school-age language and reading outcomes. Seventy-six infants completed resting-state fMRI scans during natural sleep, and their language skills, including phonological processing, were assessed behaviorally in kindergarten. Among them, 41 participants who received at least one year of reading instructions were further evaluated on their emergent word reading skills. The imaging data were analyzed to characterize the intrinsic functional organization of the emergent language network in infancy using data-driven hierarchical clustering techniques. Partial correlations were then performed to evaluate whether infant language network characteristics were associated with kindergarten-age language skills and emergent reading abilities when controlling for infant age at scan, parental education, as well as their general cognitive abilities measured at school-age. Finally, upon observed significant associations among infancy, kindergarten age and emergent reading stages, mediation analyses were applied to examine the hypothesized developmental pathway linking infant language development to word reading abilities through kindergarten-age language skills.

## 2. Materials and Methods

### 2.1 Participants

All participants were recruited as part of a longitudinal imaging project investigating the neural trajectory of long-term language and reading development since infancy (e.g., Turesky, et al., 2022; Yu et al., 2021, 2022; Zuk et al., 2021, 2022). The current study included 76 participants (33 females) who had usable resting-state fMRI scans collected in infancy (mean age = 9.3 ± 3.5 months) and underwent language skill evaluation in kindergarten (mean age = 5.8 ± 0.9 years). By the time of data analyses, a subset of 41 children (21 females) who received at least one year of formal reading instruction (i.e., emergent readers, mean age = 8.1 ± 1.1 years) were further assessed on their word reading abilities (Figure 1, Table 1). All children were born full term (gestational age ≥ 36 weeks) and exposed to English since birth. No participant had birth complications, neurological trauma, or significant developmental delays, and their infancy structural images were examined by a pediatric neuroradiologist at Boston Children’s Hospital (BCH) to ensure the absence of malignant brain features. Moreover, all subjects showed typical nonverbal IQ, demonstrated by above 80 standard scores (SS) on the Matrix Reasoning subset of the Kaufman Brief Intelligence Test-II (KBIT-II, Kaufman and Kaufman, 2004) administered at school-age. This study was approved by the Institutional Review Board at BCH and Harvard University. Informed written consent was obtained from a parent or legal guardian of each infant before participation.

**Figure 1.**
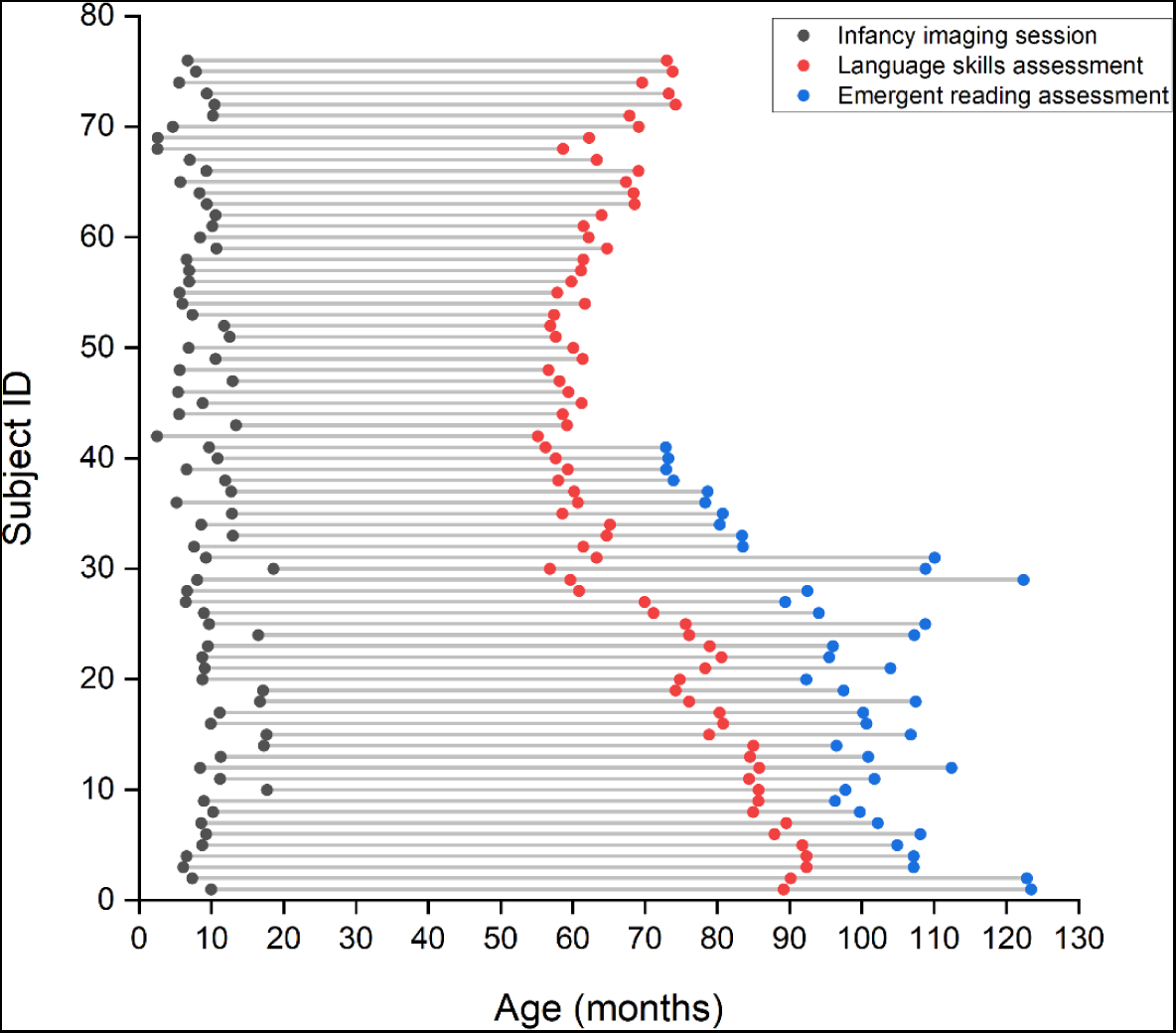
Age distribution (in months) of all included participants who completed resting-state fMRI scanning in infancy (gray dot) and were assessed in their language skills in kindergarten (red dot). A subset of children were further evaluated on their emergent word reading abilities (blue dot) after receiving at least one year of reading instructions. Dots along each horizontal line represented all available longitudinal data of one participant.

**Table 1.**
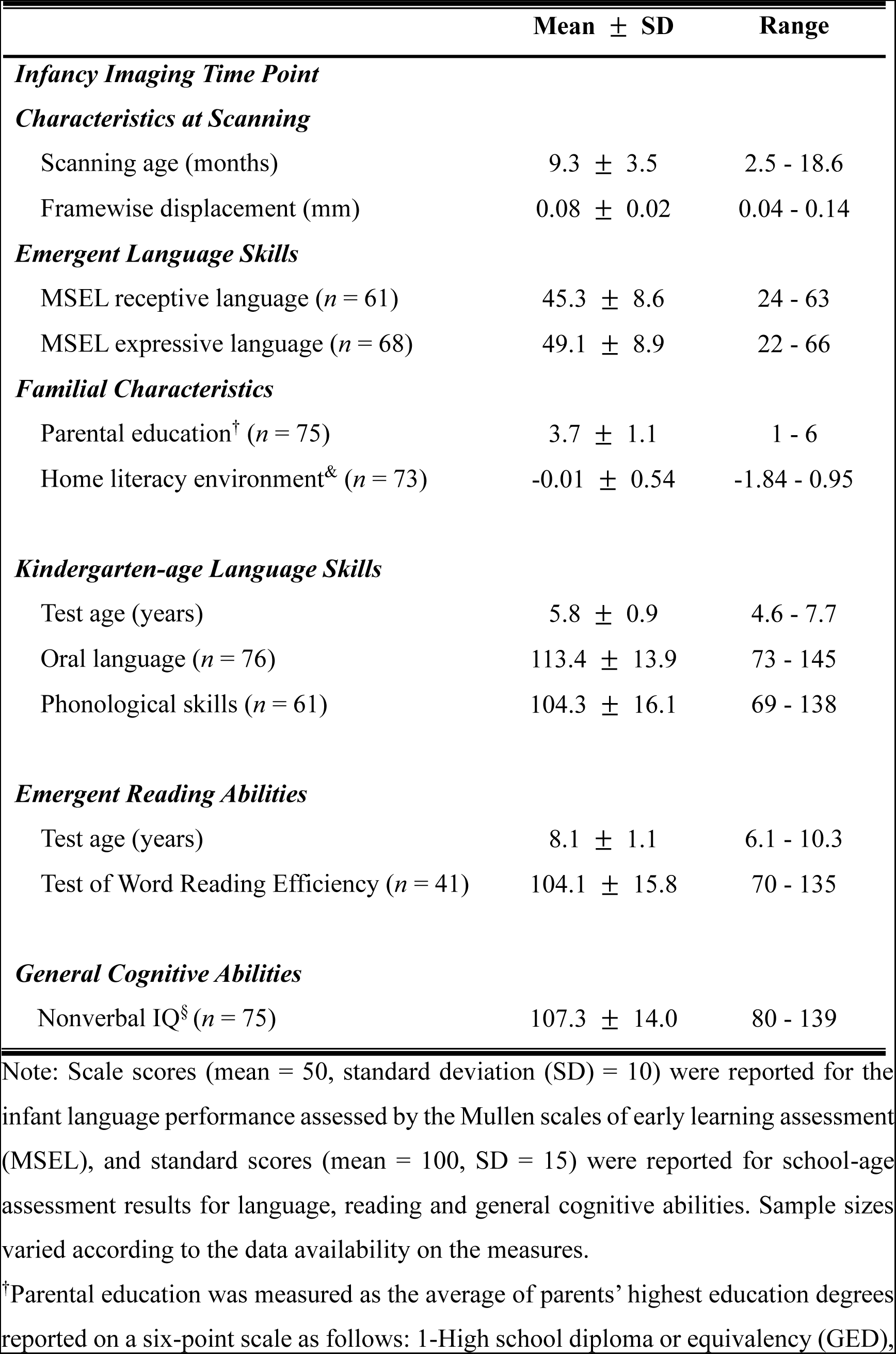
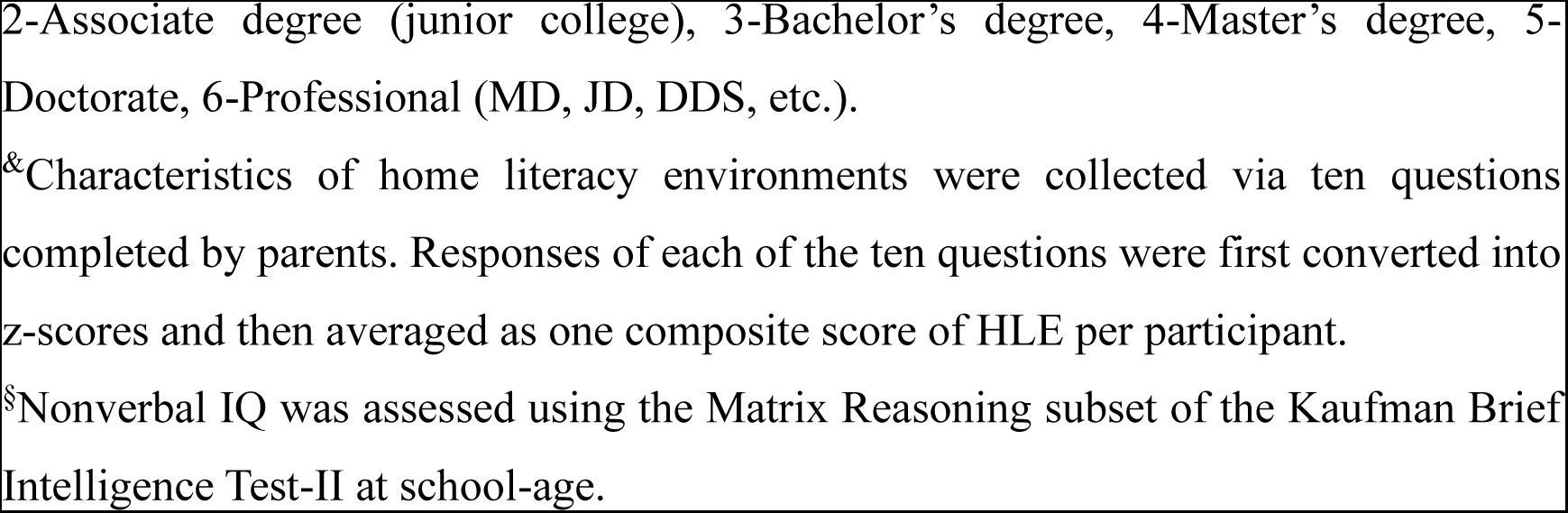
Participant characteristics at infancy, kindergarten-age and emergent reading stages.

| | Mean ± SD | Range |
| --- | --- | --- |
| ***Infancy Imaging Time Point*** |  |  |
| ***Characteristics at Scanning*** |  |  |
| Scanning age (months) | 9.3 ± 3.5 | 2.5 - 18.6 |
| Framewise displacement (mm) | 0.08 ± 0.02 | 0.04 - 0.14 |
| ***Emergent Language Skills*** |  |  |
| MSEL receptive language ($n = 61$) | 45.3 ± 8.6 | 24 - 63 |
| MSEL expressive language ($n = 68$) | 49.1 ± 8.9 | 22 - 66 |
| ***Familial Characteristics*** |  |  |
| Parental education<sup>†</sup> ($n = 75$) | 3.7 ± 1.1 | 1 - 6 |
| Home literacy environment<sup>&</sup> ($n = 73$) | -0.01 ± 0.54 | -1.84 - 0.95 |
| ***Kindergarten-age Language Skills*** |  |  |
| Test age (years) | 5.8 ± 0.9 | 4.6 - 7.7 |
| Oral language ($n = 76$) | 113.4 ± 13.9 | 73 - 145 |
| Phonological skills ($n = 61$) | 104.3 ± 16.1 | 69 - 138 |
| ***Emergent Reading Abilities*** |  |  |
| Test age (years) | 8.1 ± 1.1 | 6.1 - 10.3 |
| Test of Word Reading Efficiency ($n = 41$) | 104.1 ± 15.8 | 70 - 135 |
| ***General Cognitive Abilities*** |  |  |
| Nonverbal IQ<sup>§</sup> ($n = 75$) | 107.3 ± 14.0 | 80 - 139 |

Note: Scale scores (mean = 50, standard deviation (SD) = 10) were reported for the infant language performance assessed by the Mullen scales of early learning assessment (MSEL), and standard scores (mean = 100, SD = 15) were reported for school-age assessment results for language, reading and general cognitive abilities. Sample sizes varied according to the data availability on the measures.
<sup>†</sup>Parental education was measured as the average of parents' highest education degrees reported on a six-point scale as follows: 1-High school diploma or equivalency (GED),
2-Associate degree (junior college), 3-Bachelor's degree, 4-Master's degree, 5-Doctorate, 6-Professional (MD, JD, DDS, etc.).
&Characteristics of home literacy environments were collected via ten questions completed by parents. Responses of each of the ten questions were first converted into z-scores and then averaged as one composite score of HLE per participant.
§Nonverbal IQ was assessed using the Matrix Reasoning subset of the Kaufman Brief Intelligence Test-II at school-age.

### 2.2 Psychometric and familial characterization at infancy and school-age time points

In infancy, participants’ emergent receptive (e.g., responsiveness to sounds and/or human voices) and expressive language (e.g., the number of vowels produced during babbling) skills were evaluated using the Mullen Scales of Early Learning assessment (MSEL, Mullen, 1995; Table 1). Scale scores with a mean of 50 and a standard deviation (SD) of 10 were acquired for both the receptive and expressive language domains. Familial characteristics relevant to long-term cognitive, language and literacy development were collected via parental questionnaires. These included parental education, measured as the average of the parents’ highest degrees of education, and home literacy environment (HLE, adapted from Denney et al., 2001), which surveyed language- and literacy-related activities at home (see a full list of survey questions in Yu et al., 2021).

Children’s language skills were assessed again in kindergarten (at the beginning and/or end of), this time using the Oral Language (OL) cluster and the Phonological Processing subtest of the Woodcock-Johnson IV Tests (WJ-IV; Schrank and Wendling, 2015). The OL cluster is comprised of two subtests, which were picture vocabulary, measuring children’s ability to identify pictured objects, and oral listening comprehension, requiring children to complete audio-recorded passages with appropriate words. The Phonological Processing subtest included 1) Word Access, which required participants to provide a word that has a specific phonemic element in a specific location; 2) Word Fluency, which measured the child’s ability to name as many words as possible beginning with a specified sound in 1 minute; and 3) Substitution, which asked participants to substitute a word part to create a new word. The composite SS for the OL cluster and the Phonological Processing subtest were calculated automatically by the WJ-IV online scoring system.

Children who received at least one year of reading instruction (i.e., beginning at the end of kindergarten or later) were tested on emergent reading abilities using the Test of Word Reading Efficiency (TOWRE-2; Torgesen et al., 2012). As a timed measure of reading efficiency, TOWRE used both printed words (in the Sight Word Efficiency (SWE) subtest) and phonemically regular nonwords (in the Phonemic Decoding Efficiency (PDE) subtest) to measure the ability to pronounce words and nonwords accurately and fluently, respectively. Reading performance was obtained by averaging SS of the two subtests for each child. For children who completed the same assessments multiple times, only scores from their latest assessment were used in statistical analyses. To ensure that associations between kindergarten language skills and subsequent reading abilities could be evaluated prospectively, language skills were always assessed in the time point earlier than that for reading assessments (assessment interval = 22.6 ± 11.2 months, range: 10.6-62.7 months, Figure 1). Nevertheless, we have demonstrated that the obtained result patterns could be replicated using the mean scores of each assessment across all the available time points (Supplementary Table 1).

### 2.3 Image acquisition

Brain images of each infant were acquired during natural sleep on a Siemens 3T Trio MRI scanner with a 32-channel adult head coil (Raschle et al., 2012). Structural images were obtained using a T1 multi-echo MP-RAGE sequence with prospective motion correction (imaging parameters: TR = 2270 ms, TE1,2,3,4 = [1.66, 3.48, 5.3, 7.12] ms, TI = 1450 ms, flip angle = 7°, field of view = 220 mm^2^, voxel size = 1.1 × 1.1 × 1.0 mm^3^, number of slices = 176). An 8-minute EPI BOLD sequence was utilized to collect resting-state fMRI images. For images collected before 2017 (n = 43), the following acquisition parameters were used: TR = 3000 ms, TE = 30 ms, flip angle = 60°, voxel size = 3 × 3 × 3 mm^3^. A simultaneous multi-slice (SMS) imaging technique with a shorter TR (950 ms) was utilized for resting-state fMRI acquisition in 2017 and after (n = 33) with the other parameters kept the same. Post-processing data harmonization was performed (see details in 2.5.2) in order to accommodate the potential differences induced by different TRs.

### 2.4 Imaging data preprocessing

An infant-appropriate preprocessing pipeline (Yu et al., 2021, 2022) was applied in the current study. Functional MRI images were first corrected for slice timing using FSL/SLICETIMER (separately on each SMS slice group for the SMS-acquired data, Viessmann et al., 2019). Rigid body alignment for motion correction and estimation of motion regressors were performed using FSL/MCFLIRT. Framewise displacement (FD) was estimated based on head movement estimates (Power et al., 2012) using an in-house script (https://github.com/xiyu-bnu/infant_restingstate_prediction). Volumes with FD > 0.3 mm were considered outliers. One preceding and two subsequent artifactual frames were also removed to minimize the potential spreading effect. An average of 2.6% (SD = 0.04, range: 0% ∼ 17.6%) outlier scans were identified and removed from each participant dataset, leaving at least 5-minute usable volumes for every infant included. All fMRI data were then spatially normalized to the UNC 1-year-old infant template (Shi et al., 2011) via the corresponding structural image using affine transformations (FSL/FLIRT). Linear regression was subsequently performed with six continuous motion regressors, mean CSF and WM signals, and motion outliers as nuisance regressors. Images were further temporally band-pass filtered (0.01-0.1 Hz) and spatially smoothed (Gaussian Filtered, FWHM = 6 mm). Finally, to keep temporal resolution consistent across the whole sample, we resampled images that were acquired with the multi-slice sequence to TR = 3000ms.

### 2.5 Functional connectivity analyses of the emergent language network in infancy

#### 2.5.1 Seed selection

Seed regions constituting the language network were defined based on the language parcels developed by Fedorenko and colleagues. These parcels were generated based on the group-level activation result of the *Sentences* > *Nonwords* contrast in a reading task of 220 adult participants (Fedorenko et al., 2020; Mineroff et al., 2018; available at https://evlab.mit.edu/funcloc/), and were replicated using the *Intact* > *Degraded auditory passages* contrast in a listening comprehension task (Scott et al., 2017), indicating modality-independent language processing in these parcels. Although a left-lateralized language network is typically observed in adults, previous studies have suggested bilateral neural engagement for language processing at early developmental stage. Both young children (4-6 years old) and infants have shown bilateral activations in the frontal and temporal language regions during speech perception (May et al., 2018; Olulade et al., 2020; Perani et al., 2011). Moreover, significant interhemispheric connections have been observed among homotopic language regions of infants (Scheinost et al., 2021; Smyser et al., 2016), which were further linked to their subsequent language abilities at 4 years old (Emerson et al., 2016). Therefore, twelve bilateral regions of interest (ROIs) were included in the current study, which are the inferior frontal gyri *pars triangularis* and *pars opercularis* (termed as IFG for short), inferior frontal gyri *pars orbitalis* (IFGorb), middle frontal gyri (MFG), anterior (AntTemp) and posterior (PostTemp) part of middle temporal gyri, and angular gyri (AngG). It has recently been demonstrated that this language network exhibits selective activation for high-level linguistic processes in adults (Fedorenko et al., 2011) across 45 languages and 12 language families (Malik-Moraleda et al., 2022), thus representing a universal neural template for language processing. All ROIs were further refined using a probabilistic atlas built on the convergence (i.e., percentage of participants) of language-specific activation across 806 adults (LanA; Lipkin et al., 2022). Specifically, a region-level threshold of 25%, representing the top quarter of voxels with most consistent inter-participant responses to language processing within each region, was applied to enhance each ROI’s functional specificity for language processing. Finally, all ROIs defined in adult MNI space, were transformed to the UNC 1-year-old infant template space (Shi et al., 2011, Fig. 2A) using Advanced Normalization Tools (ANTs, Avants et al., 2009, also see Kamps et al., 2020; Lesinger et al., 2023; Truzzi and Cusack, 2023 for the application of adult functional network regions in the infant network analyses).

**Figure 2.**
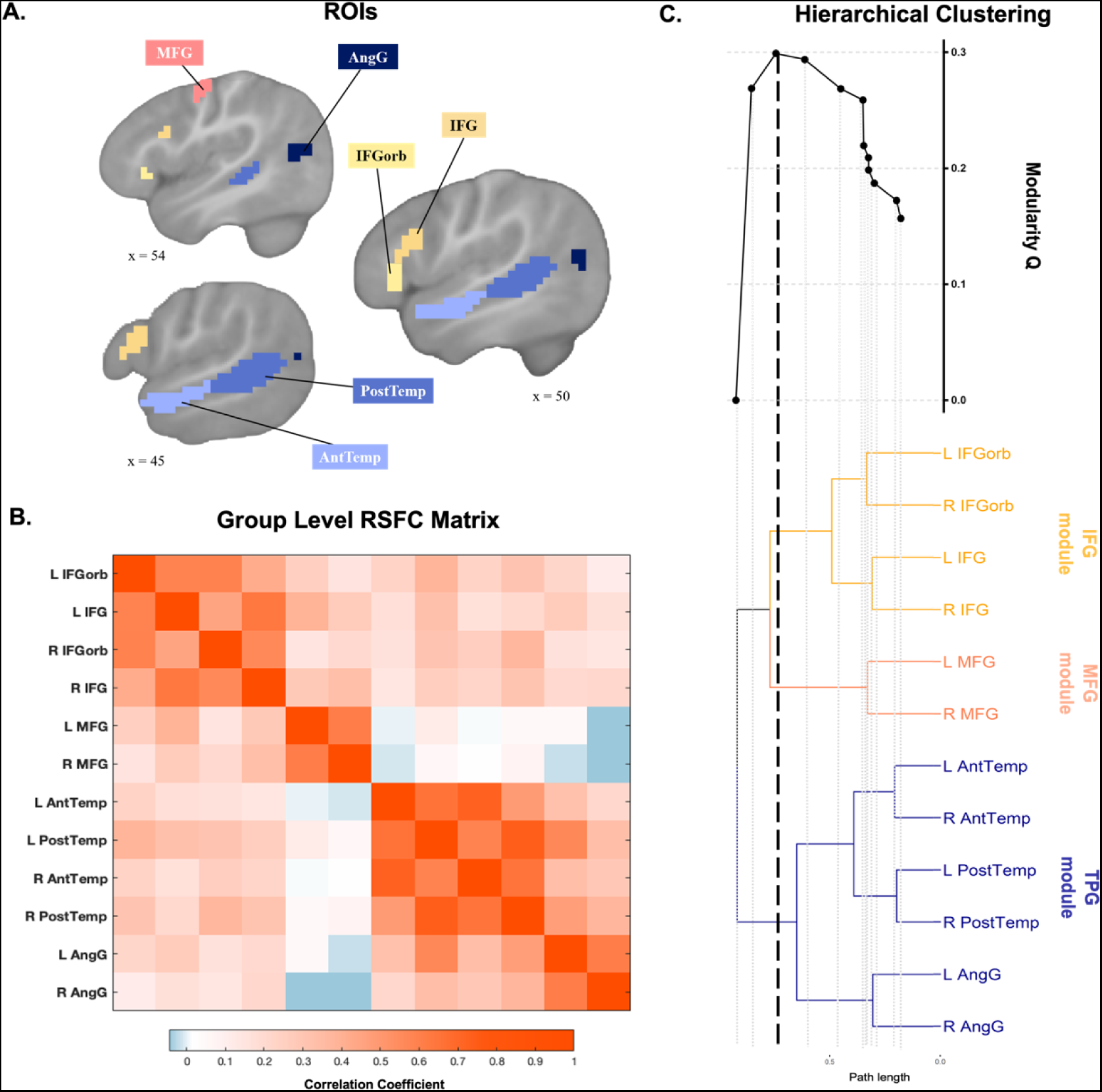
Functional connectivity and modular organization of the emergent language network in infancy. (A) Bilateral ROIs of the language network were derived from the LanA probabilistic functional atlas (region-level threshold: 25%) and transformed to the UNC 1-year-old infant template (only ROIs in left hemisphere were illustrated here). IFG: inferior frontal gyrus *pars triangularis* and *pars opercularis*; IFGorb: IFG *pars orbitalis*; MFG: middle frontal gyrus; AntTemp: anterior part of middle temporal gyrus; PostTemp: posterior part of middle temporal gyrus; AngG: angular gyrus. (B) Group level functional connectivity matrix of the emergent language network in infancy. (C) Hierarchical clustering results revealed an optimal three-module structure for the emergent language network in infancy. A binary tree created from hierarchical clustering is shown in the lower panel, which indicates the number and components of the divided clusters as the path length increases. In the upper panel, modularity values were plotted for all partition levels licensed by the binary tree. Each point on the modularity plot corresponded to a partition level indicated by the corresponding vertical line where the modular structure was defined based on branches on the right of the line. A tripartite modular architecture was indicated by a black dashed vertical line that had the highest modularity Q scores (Q = 0.30) among all possible partitions (indicated by gray dashed vertical lines) generated by the tree structure. IFG module: inferior frontal module; MFG module: middle frontal module; TPG: temporal and parietal module.

#### 2.5.2 Modularity analyses

Given the modular structure of the infant language network reported in the previous study (King et al., 2021), the data-driven clustering techniques were applied in the current study to identify potential clusters (i.e., modules) of nodes based on the infant FC patterns of the language network (see similar approaches in Blank et al., 2014; Mineroff et al., 2018). Specifically, the FC patterns were first computed for each infant by performing Pearson correlations between the mean BOLD time series of each seed region. To accommodate potential influences of different TRs on FC estimates, correlation matrices for all infants were harmonized using the ComBatHarmonization approach (https://github.com/Jfortin1/ComBatHarmonization), a method that has been reliably applied to integrate multi-site neuroimaging data (Fortin et al., 2017, 2018; Yu et al., 2018). The Fisher-Z scores derived from the harmonized correlation matrices were then averaged across all infants and transformed back to *r* values, resulting in a group level FC matrix of the infant language network (Fig. 2B). For the clustering analyses, a group level distance matrix was created by subtracting the group level FC matrix from 1. This distance matrix was submitted into the hierarchical clustering analyses using the agnes function in the R package *cluster* (Maechler et al., 2012). These analyses produced a binary tree structure, which indicated how the whole network could be divided into different clusters and which regions constitute each cluster as the path length increases (Fig. 2C). Clustering was generated based on the average linkage, where the distance between two clusters was defined as the average distances between all node pairs in the two cluster. Then, to determine the optimal partition level in the resulting binary tree structure (i.e., the “ideal” modular structure), a modularity-optimization method was applied (Newman and Girvan, 2004) using the Modularity Calculation program available in Radatools (https://deim.urv.cat/~sergio.gomez/radatools.php). To do this, all 12 possible modularity structures were delineated by increasing the path length in the binary tree structure one step at a time, ranging from a twelve-module structure with each module consisting of one single ROI to a one-module structure with all 12 ROIs as one module. A modularity Q score was computed for each of these partition results by comparing the functional correlations of the cluster structure with the expected values under a null model (Bazzi et al., 2016). Here, a uniform null model was used, which is shown to be appropriate for analyses on correlation matrices of the imaging data (Kenett et al., 2020). The optimal modular structure of the emergent language network in infancy was then selected based on the highest Q score that indicated the best clustering solution. Finally, FC of each module within the emergent language network was calculated for every infant by averaging the harmonized FC of all paths within one module, and these values were converted into Fisher-Z values for subsequent analyses.

### 2.6 Longitudinal analyses between infancy and school-age time points

#### 2.6.1 Association analyses of behavioral outcomes measured at all time points

Infants’ language scores (i.e., MSEL receptive and expressive language) were first Pearson correlated with their kindergarten-age language performance (i.e., oral language and phonological skills) and emergent reading abilities to evaluate the long-term association effects of infants’ language behaviors. Then, to estimate the concurrent associations among kindergarten-age language abilities, as well as their longitudinal relationships with emergent word reading skills, Pearson correlations were further conducted between psychometric results obtained at the school-age timepoints.

#### 2.6.2 Association analyses between infant language network characteristics and language and reading skills measured concurrently and longitudinally

Partial correlations were conducted to examine associations of infant language network characteristics with emergent language skills in infancy, as well as with school-age language and word reading abilities. Covariates included in these partial correlation analyses were selected through a two-step procedure, in order to balance the trade-off between statistical power and estimation accuracy. Specifically, infant age and head movement (i.e., mean FD values) during scanning were first selected as potential confounding factors due to their reported influences on the functional connectivity measures (e.g., Filippi et al., 2021; Satterthwaite et al., 2012; Van Dijk et al., 2012). Non-verbal IQ, parental education and HLE were also considered based on their reported relationships with language and reading development (e.g., nonverbal IQ: Lonigan et al., 2009; Stanovich et al., 1984; parental education: Friend et al. 2008; Lyytinen et al., 1998; HLE: Burgess et al., 2002; Sénéchal and LeFevre, 2014). To account for missing data in potential covariates (one nonverbal IQ, one parental education, and three HLE estimates), multiple imputation procedures implemented in IBM SPSS 26.0 were applied with imputed values averaged across five imputation samples. No data imputation was performed for behavioral outcomes of interests. The effects of these potential confounding factors on infant language network characteristics and behavioral measures at the three time points were empirically examined using correlation analyses. We observed significant correlations of children’s non-verbal IQ (*r* = 0.46, *p_corrected_* < 0.001) and parental education (*r* = 0.39, *p_corrected_* = 0.007) with their kindergarten-age oral language scores, as well as significant impacts of infant age at scan on infant FC of the language network (IFG module: *r* = −0.37, *p_corrected_* = 0.007; TPG module: *r* = 0.39, *p_corrected_* = 0.007). No other significant partial correlations were found (all *p_corrected_* > 0.05, Supplementary Table 2). Based on these results, infant age at scan, non-verbal IQ and parental education were included as covariates in the partial correlations between infant FC and verbal skills measured concurrently and longitudinally (also see Hardi et al., 2023; Smith et al., 2018; Song et al., 2015; Vanes et al., 2023 for the inclusion of control variables measured at multiple time points in their longitudinal investigations).

To investigate concurrent associations between neural and behavioral correlates of language development in infancy, partial correlation analyses were conducted between FC of each module within the infant language network and scale scores of MSEL receptive and expressive language. Subsequently, partial correlations were performed between infant FC of each language network module and standard scores of kindergarten-age phonological processing and oral language tests, to assess longitudinal relationships between infant language network characteristics and long-term language development. Finally, the same partial correlations were performed between infant FC of each module and word reading abilities of participants who received at least one year of reading instructions, to assess the prospective associations between the emergent language network in infancy and long-term reading achievement.

#### 2.6.3 Mediation analyses for longitudinal relationships between infant language network characteristics and emergent word reading abilities via kindergarten-age language skills

If any of the language skills measured in kindergarten showed significant associations with both infant FC and emergent reading abilities, the potential mediation role of the identified kindergarten-age skill was examined. To do this, a sub-group of participants that had relevant data at all time-points were selected (Figure 1). Mediation analyses were conducted via structural equation modeling, with infant age at scan, non-verbal IQ and parental education included as covariates. The statistical significance of the indirect effect (i.e., of the mediator) was computed using an estimate of the delta method standard error and compared against zero using a Z test, following Sobel’s approach (Sobel, 1982). Moreover, the effect size of the observed mediation was further estimated using Kappa-squared (𝜅^2^, Preacher and Kelley, 2011), which represents the proportion of the observed indirect effect to the maximum possible indirect effect and can be interpreted based on Cohen’s (1988) guidelines for magnitudes of effect sizes (i.e., 𝜅^2^ of 0.01, 0.09, 0.25 indicated small, medium, and large effect sizes respectively). All statistical analyses were performed using JASP 0.17.1 (Love et al., 2019), and results were adjusted for multiple comparisons using FDR corrections.

#### 2.6.4 Power analyses

Power analyses were conducted to ensure sufficient sample size to detect potential longitudinal association effects. For a power level equal to 80% (alpha = 0.05) and predicted positive correlation effects of 0.4 based on previous research examining longitudinal relationships of language measures in early childhood (Duff et al., 2015; Silvén et al., 2002; Yu et al., 2021; Zuk et al., 2021), a sample size of 37 participants was required for longitudinal association analyses between infant and school-age measures. Similarly, for correlation analyses between language skills and reading abilities with a predicted effect size of 0.6 (Melby-Lervåg et al., 2012; Torgesen et al., 1994), a sample size of 15 subjects was needed to achieve a power level of 80%. Additionally, Monte Carlo power analyses were conducted for the mediation analyses. Based on predicted correlation effects of 0.6 between language skills and reading abilities, and 0.4 for longitudinal associations of infant measures, a sample size of 46 subjects was determined for a power level of 80%.

#### 2.6.5 Follow-up analysis 1: examining longitudinal associations across infant age at scan

Infants undergo significant brain and language development within the first years of life (Gilmore, 2018; Kuhl, 2004). Therefore, additional analyses were conducted to estimate whether the observed relationships between infant FC of the IFG module and school-age phonological and word reading abilities remained consistent across infant age at scan. For this aim, we first replicated the partial correlation analyses within subsets of infants whose images were collected within the first year of life (for the phonological association: *n* = 51, mean age = 8.4 ± 2.2 months; for the word reading association: *n* = 31, mean age = 8.8 ± 1.7 months). This period is critical for the development of speech perception specific to the native language, which typically matures around 12 months old (Kuhl, 2004). Additionally, we generated another 10 subsamples of infants with increasing ages for each longitudinal association by sliding the age window along the dimension of infant age at scan until reaching the oldest age. The same partial correlations were run in each subsample to assess whether the observed longitudinal associations based on all data of infants remained significant across these subsamples covering variable portions of the entire range of infant ages.

#### 2.6.6 Follow-up analysis 2: examining longitudinal associations with phonological-related brain regions

The language network ROIs employed in the main analyses were derived by contrasting real words and sentences (task) with the nonwords and acoustically-degraded speech (i.e., control, Fedorenko et al., 2010; Scott et al., 2017). Since the word/speech-like stimuli in the control condition might entail sublexical (phoneme and syllable) processing, phonological-related processes could be largely reduced in the current language network (however, see Regev et al., 2024). Indeed, the activation-based meta-analysis map generated from the Neurosynth database comprising a total of 14,371 fMRI studies (version 0.7 released July, 2018, Yarkoni et al., 2011, term: “phonological”) revealed widespread activations for phonological processing. This map encompassed not only the classic fronto-temporal language network areas that were applied in the main analyses, but also regions in the left inferior parietal lobule (LIPL), left supramarginal gyrus (LSMG), middle and posterior parts of left superior temporal gyrus (LSTG) and the middle part of right superior temporal gyrus (RSTG), which is consistent with previous studies involving school-age children and adults (Pollack and Ashby, 2018; Tan et al., 2005; Vigneau et al., 2006, 2011). Therefore, to directly examine potential relationships between infant FC of phonological-related regions and long-term language and word reading development, four additional regions of interests (ROIs) were generated by overlaying the phonological-related meta-analysis map with each of the anatomical regions (i.e., LIPL, LSMG, LSTG and RSTG) defined by automated anatomical labeling atlas (AAL, Tzourio-Mazoyer et al., 2002). These ROIs were then transformed from adult MNI space to the UNC 1-year-old infant template space (Shi et al., 2011, Figure S1A) using ANTs. Two lines of analyses were performed. First, we tested whether infant FC between any phonological-related ROI and every module within the original language network employed in the main analyses were correlated with school-age language and word reading performance of the same children. To achieve this, partial correlations with the same covariates as applied in the main analyses were conducted. Second, we combined these four new ROIs with the original language network regions into an extended language network. Similar to the procedures in the main analyses, modularity analyses were performed to identify potential modules within this extended language network, and longitudinal associations between infant FC of each module and long-term behavioral outcomes were evaluated using partial correlations.

## 3 Results

### 3.1 Intrinsic modular architecture of the emergent language network in infancy

An overarching tripartite modular architecture (Fig. 2C) was identified in the tree structure obtained from hierarchical clustering, which had the highest modularity value (Q = 0.3) compared to all other partitions. Accordingly, the emergent language network in infancy was optimally characterized by an internal tripartite modular structure, which included an inferior frontal module involving bilateral IFG and IFGorb gyri (i.e., IFG module), a middle frontal module comprised of bilateral MFG (i.e., MFG module), and a temporal and parietal module containing anterior and posterior temporal and angular gyri bilaterally (i.e., TPG module). The same modular results were also obtained based on the thresholded group level FC matrix that only included positive functional connections (Supplementary Table 3), speaking to the robustness of the observed tripartite modular structure for the emergent language network in infancy.

### 3.2 Association results of behavioral outcomes measured at infancy and school-age time points

The correlation results of behavioral performance were presented in Table 2 (see similar results with testing ages as additional covariates in Supplementary Table 4). Specifically, infants’ emergent language skills measured as MSEL receptive and expressive language scores correlated with each other (*r* = 0.44, *p*_corrected_ = 0.002); however, neither were significantly correlated with any language or reading measure assessed at school-age (all *p* > 0.1).

**Table 2.**
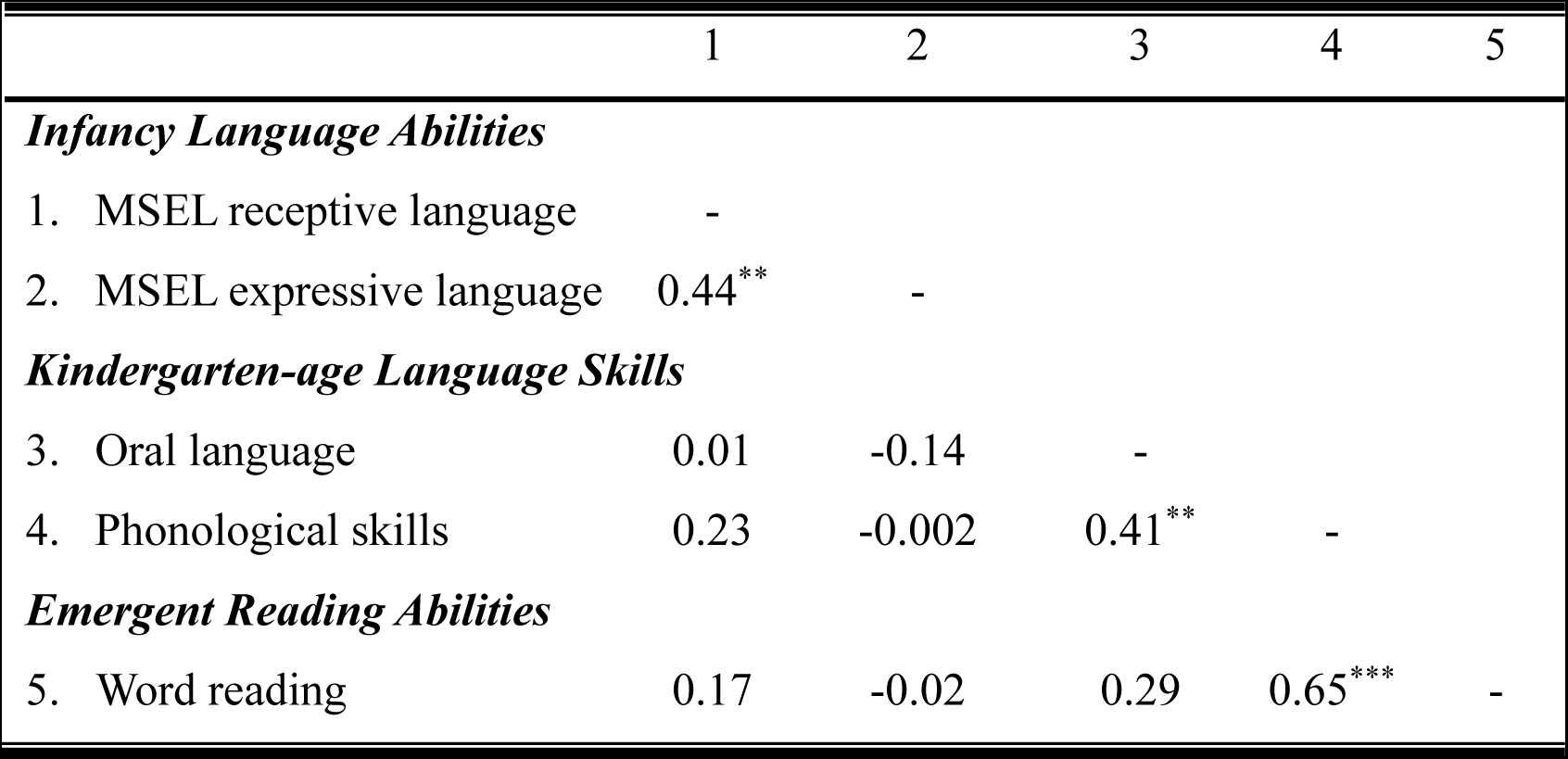
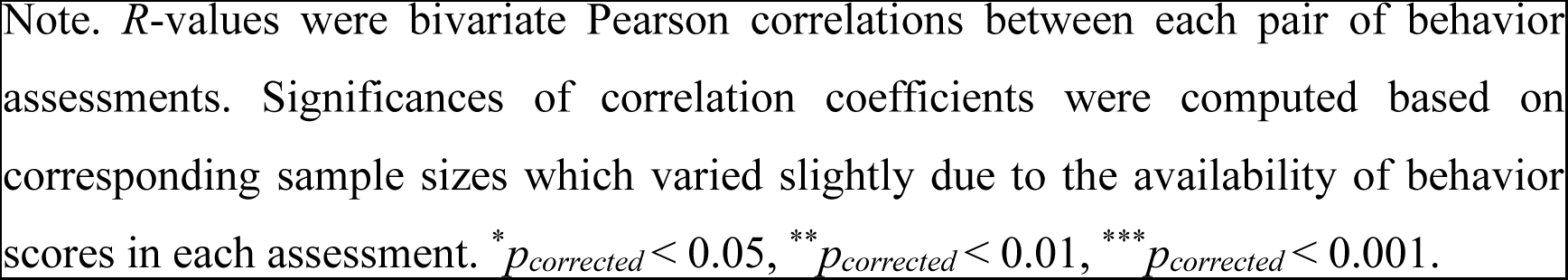
Longitudinal correlation results of behavioral outcomes between infancy and school-age time points.

|  | 1 | 2 | 3 | 4 | 5 |
| --- | --- | --- | --- | --- | --- |
| ***Infancy Language Abilities*** |  |  |  |  |  |
| 1. MSEL receptive language | - |  |  |  |  |
| 2. MSEL expressive language | 0.44** | - |  |  |  |
| ***Kindergarten-age Language Skills*** |  |  |  |  |  |
| 3. Oral language | 0.01 | -0.14 | - |  |  |
| 4. Phonological skills | 0.23 | -0.002 | 0.41** | - |  |
| ***Emergent Reading Abilities*** |  |  |  |  |  |
| 5. Word reading | 0.17 | -0.02 | 0.29 | 0.65*** | - |

Note. $R$ -values were bivariate Pearson correlations between each pair of behavior assessments. Significances of correlation coefficients were computed based on corresponding sample sizes which varied slightly due to the availability of behavior scores in each assessment. \* $p_{corrected} < 0.05$ , \*\* $p_{corrected} < 0.01$ , \*\*\* $p_{corrected} < 0.001$ .

The language skills measured in kindergarten revealed significant positive correlations with each other (*r* = 0.41, *p*_corrected_ = 0.004). Moreover, the kindergarten-age phonological skills were also prospectively associated with the emergent word reading abilities measured approximately two years later (*r* = 0.65, *p*_corrected_ < 0.001).

### 3.3 Association results between infant language network characteristics and language and reading skills measured concurrently and longitudinally

Partial correlation results of infant language network characteristics with concurrent and longitudinal behavioral measures were shown in the Supplementary Table 5. Specifically, no concurrent associations were observed between infant FC of any language network module and MSEL receptive and expressive language performance (all *p* > 0.2, Supplementary Table 5). Regarding the longitudinal associations with kindergarten-age language skills, positive partial correlations were demonstrated between FC of the IFG module in infancy and phonological skills assessed at kindergarten (*r* = 0.38, *p_corrected_* = 0.018; Fig. 3A). These associations were observed when controlling for infant age at scan, their nonverbal IQ as well as parental education. Moreover, the results were significant across the age ranges of phonological assessment (Supplementary analyses). No significant partial correlations were found for kindergarten-age oral language performance, nor the other two modules (i.e., MFG and TPG modules, all *p* > 0.2, Supplementary Table 5). Finally, for long-term reading development, partial correlations revealed significant longitudinal associations between FC of the IFG module and emergent word reading abilities (*r* = 0.43, *p_corrected_* = 0.021; Fig. 3A), a pattern that remained robust across the age ranges of reading assessment (Supplementary analyses). Infant FC of the MFG and TPG modules did not show significant partial correlations with children’s emergent reading abilities (all *p* > 0.5, Supplementary Table 5).

**Figure 3.**
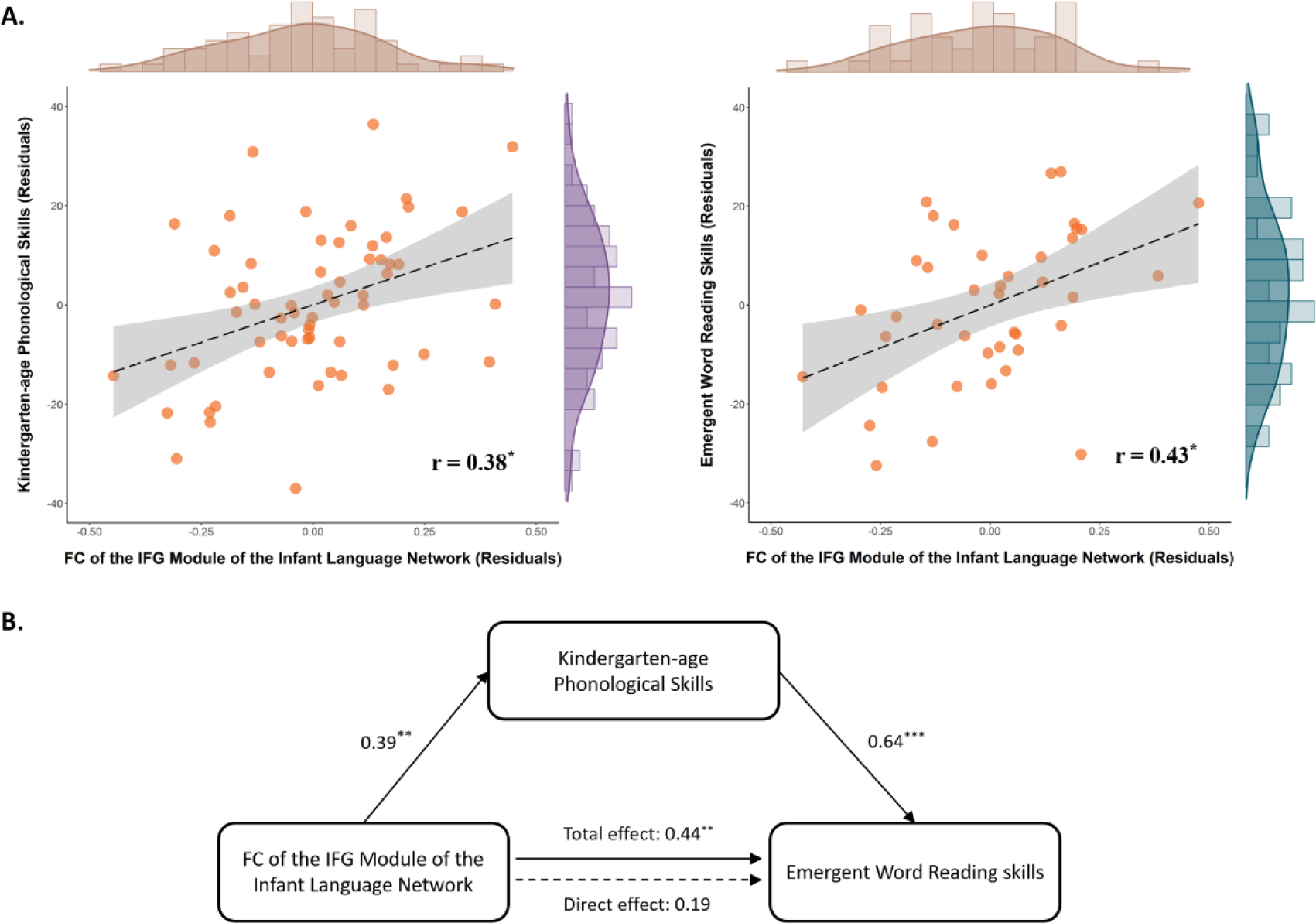
Prospective associations between infant language network characteristics and school-age language and reading outcomes. (A) Scatterplots showed significant associations between FC of the IFG module within the emergent language network in infancy and kindergarten-age phonological skills (*n* = 61, left), as well as between the infant FC of the IFG module and the emergent word reading skills (*n* = 41, right). Residual values controlling for infant age, nonverbal IQ and parental education were shown. Histogram and density estimation plots along the perimeter showed distributions of the FC of the IFG module (x-axis) and behavioral outcomes (y-axis). \**p_corrected_* < 0.05 (B) The mediation model illustrated a significant mediation role of kindergarten-age phonological skills on the relationship between the infant FC of the IFG module and emergent word reading skills (*n* = 37). Standardized effect estimates were shown, and the dotted arrow represented the insignificant direct effect after accounting for the mediation effect of the phonological skills. \*\**p* < 0.01, \*\*\**p* < 0.001

### 3.4 Mediation results for longitudinal relationships between infant language network characteristics and emergent word reading abilities via kindergarten-age phonological skills

Given the significant associations of kindergarten-age phonological skills with infant FC of the IFG module within the emergent language network and subsequent word reading achievement, the potential mediation role of phonological skill was further examined in participants whose kindergarten-age phonological and emergent reading performance were both available (*n* = 37; Fig. 3B). Note that even though the current sample was determined to be insufficient by Monte Carlo power analyses, exploratory mediation analyses were still conducted since significant longitudinal associations between the independent variable (infant FC), the mediator (kindergarten-age phonological skills) and the outcome variable (emergent reading abilities) were evident in subject samples with sufficient power. The total effect of infant FC of the IFG module on emergent reading abilities was significant based on data of this sample (total effect = 0.44, *standard error (SE)* = 0.15, 95% CI = [0.14, 0.74]), replicating the main result. Notably, an indirect effect of infant FC on emergent reading abilities via kindergarten-age phonological skills was also found (indirect effect = 0.25, *SE* = 0.11, 95% CI = [0.04, 0.46], indirect/total effect = 56%) and further supported by a Sobel test (*z* = 2.34, *p* = 0.019), while the direct effect from infant FC to emergent reading skills after accounting for the indirect pathway was not significant (direct effect = 0.19, *SE* = 0.14, 95% CI = [-0.08, 0.46]). This indirect effect indicated that kindergarten-age phonological skills mediated 56% (indirect/total effect = 56%) of the association between infant FC of the emergent language network and subsequent word reading abilities, which constituted a medium to large mediation effect based on the measure of kappa-squared (𝜅^2^ = 0.25, 95% CI [0.03, 0.47], Preacher and Kelley, 2011).

It is important to note that the result patterns obtained based on ROIs derived from the probabilistic atlas (LanA) using a region-level threshold of 25% (i.e., the threshold used to refine the ROIs used in the preceding analyses) were also evident when other thresholds (50%, 75%, 100%; Supplementary Table 6) were applied, suggesting the consistency of results across different functional specificity levels of the language network. Moreover, similar results could also be observed when using scores of the SWE and PDE subtests of the TOWRE assessment, respectively (Supplementary Table 7), further speaking to the general word reading abilities examined in the current study.

### 3.5 Longitudinal association results across infant age at scan

For the longitudinal associations between infant FC of the IFG module and kindergarten-age phonological skills, significant partial correlations were observed in the subsample of infants whose images were collected within the first year of life (*r* = 0.34, *p_corrected_* = 0.020), as well as all subsequent subsamples with increasing infant ages (all *p_corrected_* < 0.05, Supplementary Table 8). Similarly, for longitudinal associations with emergent word reading skills, significant partial correlations were demonstrated based on imaging data acquired within the first year of life (*r* = 0.40, *p_corrected_* = 0.039), as well as data from subsequent ten subsamples with increasing infant age (all *p_corrected_* < 0.05, Supplementary Table 9).

### 3.6 Longitudinal association results with phonological-related brain regions

Infant FC between each phonological-related ROI and each of the three modules within the original language network were first subjected to partial correlation analyses with the same covariates described above. These analyses revealed no significant associations with school-age oral language, phonological processing and word reading measures (all *p_corrected_* > 0.1, Supplementary Table 10). Modularity analyses with the extended language network revealed a five-module structure (Figure S1B, Supplementary Table 11). The IFG and MFG modules identified based on the original language network were retained. The two bilateral angular gyri (AngG) regions were separated from the original TPG module, forming a third module (i.e., AngG module). The two newly added phonological-related temporal regions (i.e., L STG and R STG), together with temporal regions (i.e., bilateral anterior (AntTemp) and posterior (PostTemp) parts of middle temporal gyri) in the original language network, were incorporated into a new temporal module. The remaining two phonological-related ROIs (i.e., L IPL and L SMG) formed the fifth module, referred to as the parietal module. Partial correlation analyses between infant FC of each of the five modules and subsequent behavioral outcomes confirmed the previously observed associations between the infant FC of the IFG module and school-age phonological (*r* = 0.38, *p_corrected_* = 0.030) and word reading (*r* = 0.43, *p_corrected_* = 0.035) skills, whereas no significant results were found for the other four modules (all *p* > 0.1, Supplementary Table 12).

## 4. Discussion

The present study provides the first evidence for the prospective associations between the functional organization of the emergent language network in infancy and school-age language and reading performance. Utilizing a data-driven clustering approach, we identified a tripartite modular structure for the infant language network that included inferior frontal (IFG), middle frontal (MFG), as well as temporal and parietal (TPG) modules. We further observed that infant FC of the IFG module, but not the MFG and TPG modules, was prospectively associated with kindergarten-age phonological skills and emergent word reading abilities, with the latter longitudinal relationship significantly mediated by kindergarten-age phonological skills. Moreover, the effects were observed when controlling for infant age at scan, their general cognitive abilities measured at school-age and the parental education. Overall, these findings suggest that language network characteristics in infancy may scaffold long-term language and reading development.

Intrinsic functional connections of the IFG module within the emergent language network in infancy are correlated with kindergarten-age phonological skills, highlighting the early emergence of functional circuits in the bilateral inferior frontal areas that might link to long-term phonological development. Infants show bilateral IFG activations in response to spoken language compared to non-linguistic signals (e.g., Altvater-Mackensen and Grossmann, 2018; May et al., 2018), suggesting the early involvement of bilateral IFG in emergent language processing. Moreover, specific linguistic operations have been revealed in the IFG, predominantly in the left hemisphere. For instance, the IFG has been implicated in phonetic processing in infants, as previous fNIRS and MEG studies have shown its involvement in phonemic discrimination of preterm infants prior to the complete formation of cortical layers (bilateral, Mahmoudzadeh et al., 2013) and its increased activations for native but not nonnative phonetic changes during perceptual narrowing in the second half of the first year (left, Zhao and Kuhl, 2022). Moreover, the IFG has also been associated with processing of larger units of speech sound, such as recognizing specific pattern regularities (i.e., ABB) over tri-syllabic sequences consisting of different syllables (left, Gervain et al., 2008), and decoding newly learned words from continuous speech steams (left, Minagawa et al., 2017). These task-induced functional activations imply the early functional onset of the (left) IFG in facilitating infants’ learning of the sound structural properties of one’s native language(s). Meanwhile, utilizing FC techniques, significant increases in bilateral IFG connections are evident in infancy (Scheinost et al., 2021) and prospectively associated with language development in preschoolers (Emerson et al., 2016), indicating bilateral IFG involvement in early language development, possibly through cross-hemispheric connections. The present results substantiate the putative role of bilateral IFG by relating infant FC characteristics to children’s kindergarten-age phonological skills. Importantly, this relationship was observed when participants’ general cognitive abilities and parental education were controlled, speaking to the inherent network characteristics in infancy that may underlie learning of phonological skills. Finally, it is worth noting that follow-up analyses incorporating phonological-related regions, including left inferior parietal and supramarginal gyri, as well as bilateral posterior superior temporal gyri, failed to reveal significant relationships between infant FC of these regions and subsequent behavioral measures. While null results should be interpreted with caution, the absence of longitudinal associations may suggest a later functional onset of these regions in phonological processing. Future longitudinal studies are thus needed to unravel the dynamic development of the neural correlates of phonological processing in young children. Overall, our FC-based findings complement task-based fMRI results in infancy, shedding light on the significance of the early-emerging bilateral IFG module within the infant brain for protracted language development.

Previous behavioral findings have led to a hypothesized role of early lexical development (vocabulary size) on the phonological skill acquisition (Goswami, 2007). In English, the emerging lexicons in young children are characterized by high proportion of phonologically similar entries (>80% of words have at least one phonological neighbor, Dollaghan, 1994). Therefore, as children’s vocabulary size increases, the phonologically similar neighborhoods of their lexicons also grow, which could facilitate more fine-grained analyses/representation of word sound patterns and support phonological development (Metsala and Walley, 2013; Stoel-Gammon, 2011). Consistent with this postulation, vocabulary size in late infancy (i.e., around 2 years old) is shown to be longitudinally associated with subsequent syllabic awareness skills (Duff et al., 2015; Lee, 2011; Silvén et al., 2002, 2007; Torppa et al., 2010), and commensurate vocabulary and phonological development were documented in both late talkers (Rescorla and Ratner, 1996) and lexically precocious toddlers (Smith et al., 2006). Meanwhile, infant’s left IFG responses to speech are recently shown to link to toddlerhood vocabulary growth, implying a potential role of IFG in supporting lexical development (Zhao and Kuhl, 2022). Within this context, the phonological-related FC of the IFG module could be interpreted as neural mechanisms underlying vocabulary acquisition in early childhood, which in turn foster subsequent phonological development. The IFG involvement in early lexical development is not in contrast with its proposed role in sound pattern recognition, as the latter could contribute to the sound-meaning mapping essential for vocabulary acquisition. This account could also be reconciled with the null results observed between infant FC of the IFG module and kindergarten oral language skills measured using picture vocabulary and oral comprehension tasks. While the initial word learning might rely critically on word sound identification, completion of more advanced lexical tasks would require access to the semantic knowledge represented by a distributed neural network (Binder et al., 2009; Lambon Ralph et al., 2017). Consistent with this speculation, we have shown that whole-brain FC patterns of the semantic hub, i.e., left temporal pole, in infancy is prospectively associated with school-age oral language performance (Yu et al., 2021). Given IFG’s implications in various linguistic processes during early development, future studies are warranted to unravel its functional properties in the infant brain that may contribute to the development of children’s phonological skills, vocabulary, and language abilities in general. Artificial language models capable of simultaneous learning of diverse linguistic aspects could provide valuable insights into the underlying brain mechanisms supporting the multi-level development of language abilities (Tuckute et al., 2024).

We further demonstrated that the infant FC of the IFG module within the emergent language network was positively correlated with emergent word reading abilities, and kindergarten-age phonological skills significantly mediated this relationship. Previous longitudinal studies showed that the neural activations of left IFG during oral comprehension among kindergarteners (Jasińska et al., 2021) and beginning readers (Preston et al., 2016) was predictive of subsequent reading gains, suggesting potential neural precursors of decoding abilities in kindergarteners. Meanwhile, event-related potential studies (ERP) revealed prospective associations between newborns’ neural responses to speech and their reading-related skills at age 6.5 and 2^nd^ grade (Guttorm et al., 2010; Leppänen et al., 2010). However, the limited spatial resolution of ERPs constrains the inferences on anatomical specificity. The current observation complements these findings by showing that bilateral IFG exhibits infant FC characteristics associated with school-age word reading abilities. Moreover, we identified the mediating role of kindergarten-age phonological skills in this long-term relationship. These results not only align with the well-established role of phonological skills as one of the key reading precursors (Melby-Lervåg et al., 2012; Ozernov-Palchik and Gaab, 2016; Wagner and Torgesen, 1987), but also highlight the continuous nature of reading development with phonological skills serving as an important mid-way rather than starting milestone. Overall, our results provide empirical findings of the neurobiological characteristics of the emergent language network in infancy that may underlie the development of core language skills and lay the foundation for subsequent reading acquisition.

Additionally, using a data-driving clustering approach, we identified a modular architecture of the language neural network early in life. In this architecture, homologous regions in both hemispheres of the infant brain were grouped into the same cluster. This differs from the internal architecture of the adult language network, where clustering was dominantly determined by hemisphere, grouping regions within the same hemisphere together (Blank et al., 2014; Mineroff et al., 2018). These maturational differences are in accordance with the developmental shifts from bilateral activations during speech perception in infants (Perani et al., 2011) to left-hemispheric dominance in adults (Olulade et al., 2020), suggesting postnatal development of typical left-lateralization in the language network. Furthermore, the current clustering analyses grouped the *pars triangularis*, *pars opercularis* and *pars orbitalis* subregions of IFG into the same module of the emergent language network, which are consistent with previous findings based on differently defined ROIs of IFG subregions in five-to-eight-month-old infants (King et al., 2021). However, they are different from observed clustering patterns in adults, where various IFG regions are likely to be assigned into different modules (Blank et al., 2014; Mineroff et al., 2018). Given the distinct functional roles these frontal regions play in adult language processing (Fedorenko and Blank, 2020; Hagoort, 2014), such maturational changes in clustering patterns might indicate a process of functional division and specialization within the IFG over development. Future work is imperative to unravel the specific mechanisms and the critical periods of both the hemispheric and the functional specialization within the language network over development. This knowledge may advance our understanding of the neuro-plastic nature of the infant brain that supports typical language development despite early lesions in the left-hemispheric language areas (Bates et al., 2001; Newport et al., 2022).

Finally, it is noteworthy that no significant associations were found for infant language performance, either with school-age language and reading skills or with infant language network characteristics. Previous analyses revealed mixed results on behavioral longitudinal associations of language development. Significant correlations with school-age phonological and reading development were reported for vocabulary size estimated at 2 years old using the parental questionnaire MacArthur–Bates Communicative Development Inventories (Duff et al., 2015; Lee, 2011; Torppa et al., 2010). However, vocabulary size at 1 year of age, as measured by testing infants in a laboratory setting, failed to exhibit such longitudinal correlations (Silvén et al., 2007), which is consistent with our findings. Differences in data collection methods, testing ages and longitudinal intervals might contribute to these inconsistencies. Alternatively, it is possible that early language behaviors might only indirectly affect school-age development, possibly through intermediate language skills measurable at preschool ages (Silvén et al., 2007). Meanwhile, consistent with previous literature (Bruchhage et al., 2020; Emerson et al., 2016; Ouyang et al., 2020; Sket et al., 2019), infant language network characteristics were not correlated with concurrent language behaviors in the current study, but they were related to long-term language and reading outcomes. These findings collectively may suggest a specific scaffolding role of the early-emerging language network for subsequent, rather than concurrent, development of language abilities.

There are several limitations in the current study. First, our study characterized language network characteristics based on FC patterns derived from the resting-state data of infants collected during natural sleep, precluding interpretation on the specific linguistic processes/functions these regions support. Future studies employing active language tasks are warranted to examine the functional properties of these regions (e.g., Wang et al., 2023, preprint). Second, the mediation results were derived from data with insufficient sample size (lacking 9 subjects), highlighting the need for confirmatory analyses with larger samples. Third, the current analyses only considered the language network. More work is needed to examine when and how the language network may interact with other neural networks, to facilitate the learning process of complex abilities, such as reading. Fourth, to maximize the sample size used for the longitudinal analyses, participants with longitudinal behavioral outcomes measured at variable ages, as well as two cohorts of infants with different image acquisition sequences, expressive language skills and testing ages (Supplementary Table 13) were included in the current analyses, resulting in increased variability within the dataset. Finally, it should be noted that the quantity of resting state images collected for each infant (i.e., 5 minutes) could be limited for estimating a highly-stable modularity structure (Gordon et al., 2017). To address these limitations, future studies with longer resting-state acquisition times and larger samples of children, preferably with more narrow age ranges, are needed to further evaluate the stability of long-term associations between infant language network characteristics and various behavioral measures.

## 5. Conclusions

In conclusion, employing a seven-year longitudinal study starting in infancy, the current study revealed a modular structure of the emergent language network in infancy that related to individual differences in school-age word reading and decoding achievements through a significant mediation effect of kindergarten-age phonological skills. These longitudinal effects were observed when controlling for participants’ general cognitive abilities measured at school-age and the educational level of their parents, highlighting the functional specificity of the early-emerging language network for subsequent verbal development. Overall, this work provides the first empirical evidence for the neurobiological characteristics of the early-emerging language network in infancy that may underlie children’s language development and lay a critical foundation for long-term reading acquisition.

## Data Statement

All ROIs and codes for the functional connectivity analyses in the present study are available at: https://github.com/XinyiTang16/LanNet_code. Because of the Institutional Review Board regulations at BCH, the data used in this study cannot be available on a permanent third-party archive currently. However, data sharing can be initiated through a Data Usage Agreement upon request.

## Acknowledgments

This study was supported by the Eunice Kennedy Shriver National Institute of Child Health and Human Development #R01HD065762, the National Natural Science Foundation of China #32100867 (awarded to X.Y.), the Harvard Catalyst/NIH #5UL1RR025758 (awarded to N.G.), the Charles H. Hood Foundation (awarded to N.G.), the Boston Children’s Hospital Pilot Grant (awarded to N.G.) and the National Natural Science Foundation of China #82021004 (awarded to M.X.). We sincerely thank all participating families in this longitudinal study, and also acknowledge the assistance of Yue Liu, Haojie Wen, Ziliang Zhu, and Ziyi Xiong for helpful suggestions in data analyses.

