## Supplementary materials for "Longitudinal associations between language network characteristics in the infant brain and school-age reading abilities are mediated by early-developing phonological skills"

*Supplementary analyses on the robustness of the observed longitudinal associations across specific ages of behavioral assessments for phonological and word reading abilities*

Due to the scheduling challenging for long-term longitudinal investigations, especially during pandemics, children were assessed for phonological and word reading abilities at variable ages. The large variability in ages might be a concern as children at different ages could employ different strategies for the same assessment. To address this issue, we have implemented several approaches to ensure the reliability of the observed associations across the age ranges examined at school-age time points. First, we utilized standardized scores (SS) for psychometric assessments (i.e., Receptive language and Expressive language scale of the Mullen Scales of Early Learning assessment; Oral language and Phonological processing tests of the Woodcock-Johnson IV Tests and Test of Word Reading Efficiency) in the current study. These scores are calculated based on national-wide norms, and indicate a child's performance compared to that of children in the same age group. Therefore, SS is often stable across ages. Indeed, for school-age language and word reading abilities in children with multiple performance available, we observe very high correlations between task performance based on the first and last assessment scores (phonological skills: $r = 0.93, p < 0.001$; oral language: $r = 0.94, p < 0.001$; word reading abilities: $r = 0.94$, $p < 0.001$). Moreover, none of the SS of these psychometric assessments showed significant correlations with testing ages (all $p > 0.05$). Second, we conducted post hoc analyses to re-evaluate the longitudinal associations between infant FC of the IFG module and school-age phonological and word reading abilities using behavioral scores obtained at the earliest assessment when multiple assessments were administered for the same children. Significant partial correlations were still evident (phonological processing: $r = 0.34$, $p = 0.008$; word reading abilities: $r = 0.37, p = 0.023$) when controlling for infant age at scan, non-verbal IQ and parental education, whereas no partial correlation was observed for other infant language network module or kindergarten-age oral language skills (all $p > 0.05$). Finally, we added the testing ages at the behavioral assessment timepoints as an additional control variable in partial correlations. Significant longitudinal associations between infant FC of the IFG module and school-age phonological (r = 0.38, $p_{\text{corrected}} = 0.021$) and word reading (r = 0.45, $p_{\text{corrected}} = 0.017$) performance could still be replicated, while no longitudinal associations for infant FC of other modules or kindergarten-age oral language skills were found (all $p > 0.05$, Supplementary Table 14). Altogether, these results demonstrated that the identified longitudinal relationships between infant FC and individual differences in the standardized scores of the language and reading assessments were reliable across specific testing ages.

Nevertheless, it remains possible that children at different ages relied on different yet related strategies when performing the same assessment, potentially leading to highly correlated assessment scores across different timepoints. Therefore, post hoc analyses based on subsamples of children with narrower age ranges were conducted to directly examine whether the observed associations persisted when children were at similar ages and more likely to use similar strategies. Specifically, for phonological processing, we first selected a subsample of children whose phonological skills were assessed before the age of 7 ($n$ = 47, mean age = 5.5 ± 0.6 years), as children at this age may rely more on phonological processing skills and print-sound mapping skills for sounding out words (Fox and Routh, 1975; Plaza, 2001; Rosner and Simon, 1971). Significant partial correlations were observed between infant FC of the IFG module and kindergarten-age phonological skills ($r$ = 0.42, $p_{\text{corrected}}$ = 0.014). We then generated another 13 subsamples of children ($n$ = 47) with increasing ages by sliding the window along the dimension of testing age until reaching the oldest age. The same partial correlation analyses revealed significant results in all subsamples (all $p_{\text{corrected}} < 0.05$, Supplementary Table 15), indicating that the inclusion of children that might rely on different strategies did not alter the observed longitudinal associations between infant language network characteristics and subsequent phonological skills. Applying the same analyses to the word reading performance, we first examined longitudinal associations between infant FC of the IFG module and emergent word reading abilities in a subset of children younger than 9 years old (i.e., third grade or lower, $n$ = 33, mean age = 7.8 ± 1.0 years) when phonological skills played a more important role in reading activities than in later stages (Badian, 1995, 2001; Deacon, 2012). The same analyses were then performed in another 7 subsamples of children with increasing ages generated by sliding the age window. Significant associations were present in all subsamples (all $p_{\text{corrected}} < 0.05$, Supplementary Table 16), suggesting that the observed relationships were robust across reading development.

Overall, the supplementary analyses based on the whole sample and subsamples of participants demonstrated that, despite the variability in ages of follow-up behavioral assessments, the observed longitudinal associations between infant language network characteristics and long-term phonological and word reading development were robust across the age ranges tested.

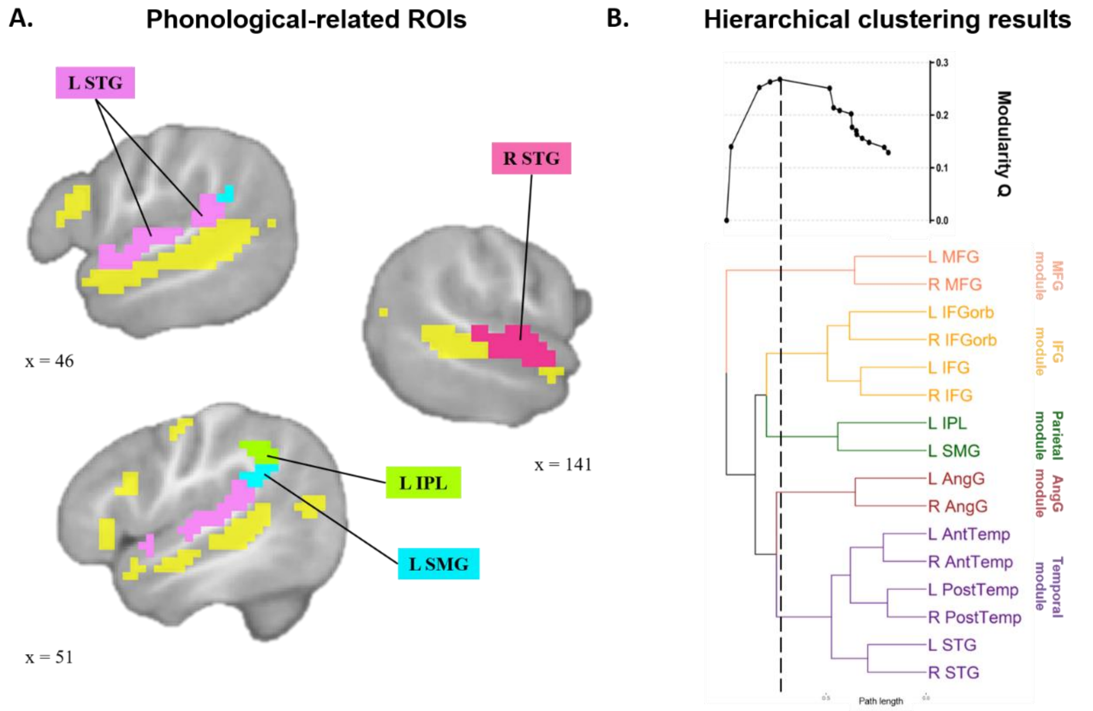


**Figure S1**. Phonological-related ROIs and modular organization of the extended language network that consisted of the original language network ROIs and four additional phonological-related ROIs. (A) Phonological-related ROIs were generated based on Neurosynth meta-analyses and transformed to the UNC 1-year-old infant template. Regions with yellow color were original language network ROIs applied in the main analyses. LIPL: left inferior parietal lobule; LSMG: left supramarginal gyrus; LSTG: left superior temporal gyrus; RSTG: right superior temporal gyrus. (B) Hierarchical clustering results revealed an optimal five-module structure for the extended language network that consisted of the original language network ROIs and four phonological-related ROIs. A binary tree created from hierarchical clustering is shown in the lower panel, which indicates the number and components of the divided clusters as the path length increases. In the upper panel, modularity values were plotted for all partition levels licensed by the binary tree. A five-module architecture was indicated by a black dashed vertical line that had the highest modularity Q scores (Q = 0.27) among all possible partitions generated by the tree structure. IFG module: inferior frontal module; MFG module: middle frontal module; AngG: angular gyri module.

**Supplementary Table 1.** Replication results based on mean scores of school-age phonological and word reading assessment performance across all available time points

| **Module** | **Partial correlations between the infant FC of the IFG module and subsequent language/reading outcomes** | | **Mediation results** | | |
|---|---|---|---|---|---|
| | **Kindergarten-age phonological processing** | **Emergent word reading skills** | **Estimates of the indirect effect and the 95% CIs** | ***z* values** | ***p* values** |
| IFG module | $r = 0.38$, $p = 0.002$ | $r = 0.39$, $p = 0.015$ | 0.27 [0.05, 0.48] | 2.45 | 0.014 |

Note:

1. Partial correlations controlled for infant age at scanning time point, nonverbal IQ and parental education.
2. The indirect effect reflected the effect of the infant FC of the IFG module on emergent word reading abilities via kindergarten-age phonological skills

**Supplementary Table 2.** Correlation results of five potential confounding factors (i.e., infant age and head movement during scanning, school-age nonverbal IQ, parental education and HLE) with infant language network characteristics and behavioral measures at all time points.

| | Infant age | Head movement | Nonverbal IQ | Parental education | HLE |
|---|---|---|---|---|---|
| ***Behavioral measures*** | | | | | |
| MSEL receptive language in infancy | -0.27 | 0.24 | -0.04 | -0.13 | -0.11 |
| MSEL expressive language in infancy | -0.24 | 0.32 | 0.02 | -0.22 | -0.16 |
| Kindergarten-age oral language | 0.09 | -0.16 | 0.46*** | 0.39** | 0.27 |
| Kindergarten-age phonological skills | 0.27 | -0.11 | 0.19 | 0.22 | 0.23 |
| Emergent word reading | 0.05 | -0.22 | 0.09 | 0.27 | 0.06 |
| ***Infant language network characteristics*** | | | | | |
| FC of IFG module | -0.37** | 0.10 | -0.12 | -0.19 | -0.11 |
| FC of MFG module | -0.02 | 0.13 | -0.20 | -0.14 | -0.16 |
| FC of TPG module | 0.39** | -0.10 | 0.11 | 0.06 | 0.19 |

Note. Correlation coefficients were bivariate Pearson correlations of potential confounding factors with behavioral measures and infant language network characteristics. IFG module: inferior frontal module; MFG module: middle frontal module; TPG module: temporal and parietal module. $^{**}p_{corrected} < 0.01$, $^{***}p_{corrected} < 0.001$.

**Supplementary Table 3.** Hierarchical clustering results based on the thresholded group level FC matrix that only included positive functional connections.

| Clustering results | | Cluster assignments of the twelve seed regions | | | | | | | | | | | | Modularity measures |
|---|---|---|---|---|---|---|---|---|---|---|---|---|---|---|
| Path length | N of Clusters | L IFGorb | L IFG | R IFGorb | R IFG | L MFG | R MFG | L AntTemp | L PostTemp | R AntTemp | R PostTemp | L AngG | R AngG | Q values |
| 0.265 | 12 | 1 | 2 | 3 | 4 | 5 | 6 | 7 | 8 | 9 | 10 | 11 | 12 | 0.16 |
| 0.281 | 11 | 1 | 2 | 3 | 4 | 5 | 6 | 7 | 8 | 9 | 8 | 10 | 11 | 0.17 |
| 0.368 | 10 | 1 | 2 | 3 | 4 | 5 | 6 | 7 | 8 | 7 | 8 | 9 | 10 | 0.19 |
| 0.389 | 9 | 1 | 2 | 3 | 2 | 4 | 5 | 6 | 7 | 6 | 7 | 8 | 9 | 0.20 |
| 0.391 | 8 | 1 | 2 | 3 | 2 | 4 | 5 | 6 | 7 | 6 | 7 | 8 | 8 | 0.21 |
| 0.408 | 7 | 1 | 2 | 3 | 2 | 4 | 4 | 5 | 6 | 5 | 6 | 7 | 7 | 0.22 |
| 0.411 | 6 | 1 | 2 | 3 | 2 | 4 | 4 | 5 | 5 | 5 | 5 | 6 | 6 | 0.26 |
| 0.496 | 5 | 1 | 2 | 1 | 2 | 3 | 3 | 4 | 4 | 4 | 4 | 5 | 5 | 0.27 |
| 0.636 | 4 | 1 | 1 | 1 | 1 | 2 | 2 | 3 | 3 | 3 | 3 | 4 | 4 | 0.29 |
| 0.747 | 3 | 1 | 1 | 1 | 1 | 2 | 2 | 3 | 3 | 3 | 3 | 3 | 3 | 0.30 |
| 0.836 | 2 | 1 | 1 | 1 | 1 | 1 | 1 | 2 | 2 | 2 | 2 | 2 | 2 | 0.27 |

Note: Red fonts indicate the modular partition with the maximum modularity Q value.

**Supplementary Table 4.** Longitudinal partial correlation results of behavioral outcomes at infancy and school-age time points controlling for test ages.

| | 1 | 2 | 3 | 4 | 5 |
|---|---|---|---|---|---|
| ***Infancy Language Abilities*** | | | | | |
| 1. MSEL receptive language | - | | | | |
| 2. MSEL expressive language | 0.43** | - | | | |
| ***Kindergarten-age Language Skills*** | | | | | |
| 3. Oral language | -0.004 | -0.15 | - | | |
| 4. Phonological skills | 0.29 | 0.09 | 0.43** | - | |
| ***Emergent Reading Abilities*** | | | | | |
| 5. Word reading | 0.30 | 0.001 | 0.39* | 0.66*** | - |

Note. Correlation coefficients were partial correlations between each pair of behavior assessments controlling for testing ages at corresponding time points. Significances of correlation coefficients were computed based on corresponding sample sizes which varied slightly due to the availability of behavior scores in each assessment. $^{*}p_{corrected} < 0.05$, $^{**}p_{corrected} < 0.01$, $^{***}p_{corrected} < 0.001$.

**Supplementary Table 5.** Partial correlation results between FC of each module within the emergent language network in infancy and behavioral outcomes controlling for infant age, nonverbal IQ, and parental education.

| | **IFG module** | **MFG module** | **TPG module** |
|---|---|---|---|
| **MSEL receptive language** | $r$ = -0.06, $p$ = 0.633 | $r$ = -0.15, $p$ = 0.264 | $r$ = -0.10, $p$ = 0.468 |
| **MSEL expressive language** | $r$ = -0.04, $p$ = 0.779 | $r$ = 0.11, $p$ = 0.393 | $r$ = 0.07, $p$ = 0.569 |
| **Phonological skills** | $r$ = 0.38*, $p$ = 0.003 | $r$ = -0.13, $p$ = 0.325 | $r$ = -0.06, $p$ = 0.655 |
| **Oral language** | $r$ = 0.11, $p$ = 0.354 | $r$ = 0.01, $p$ = 0.956 | $r$ = -0.14, $p$ = 0.241 |
| **Emergent word reading** | $r$ = 0.43*, $p$ = 0.007 | $r$ = 0.03, $p$ = 0.862 | $r$ = 0.10, $p$ = 0.542 |

Note: IFG module: inferior frontal module; MFG module: middle frontal module; TPG module: temporal and parietal module. *$p_{corrected} < 0.05$

**Supplementary Table 6.** Replication results based on ROIs derived from LanA using various region-level thresholds

| Threshold | Partial correlations between the infant FC of the IFG module and subsequent language/reading outcomes | | Mediation results | | |
|---|---|---|---|---|---|
| | **Kindergarten-age phonological skills** | **Emergent word reading skills** | **Estimates of the indirect effect and the 95% CIs** | ***z* values** | ***p* values** |
| 25% | *r* = 0.38, *p* = 0.003 | *r* = 0.43, *p* = 0.007 | 0.25 [0.04, 0.46] | 2.34 | 0.019 |
| 50% | *r* = 0.35, *p* = 0.006 | *r* = 0.44, *p* = 0.006 | 0.25 [0.04, 0.47] | 2.31 | 0.021 |
| 75% | *r* = 0.33, p = 0.011 | *r* = 0.46, *p* = 0.004 | 0.24 [0.03, 0.45] | 2.23 | 0.026 |
| 100% | *r* = 0.29, *p* = 0.028 | *r* = 0.47, *p* = 0.003 | 0.21 [0.01, 0.42] | 2.03 | 0.042 |

Note:

1. The results based on the 25% threshold were reported in the main manuscript.
2. Partial correlations controlled for infant age, nonverbal IQ, and parental education.
3. The indirect effect reflected the effect of the infant FC of the IFG module on emergent word reading abilities via kindergarten-age phonological skills.

**Supplementary Table 7.** Results based on scores of the Sight Word Efficiency (SWE) and Phonemic Decoding Efficiency (PDE) subtests of the TOWRE assessment

| **TOWRE Subtest** | **Partial correlation results** | **Mediation results** | | |
|---|---|---|---|---|
| | | **Estimates of the indirect effect and the 95% CIs** | ***z* values** | ***p* values** |
| SWE | $r = 0.40$, $p = 0.014$ | 0.20 [0.01, 0.39] | 2.06 | 0.040 |
| PDE | $r = 0.43$, $p = 0.008$ | 0.29 [0.06, 0.51] | 2.50 | 0.012 |

Note:

1. Partial correlations were performed between the infant FC of the IFG module within the emergent language network and scores of each TOWRE subtest, controlling for infant age, nonverbal IQ, and parental education.
2. The indirect effect reflected the effect of the infant FC of the IFG module on performance of each TOWRE subtest via kindergarten-age phonological skills

**Supplementary Table 8.** Partial correlation results between infant FC of the IFG module and kindergarten-age phonological skills in subsamples ($n = 51$) with increasing infant ages.

| Subsamples | Mean of infant age (months) | SD of infant age (months) | Partial correlation *r* values | *p* values |
|---|---|---|---|---|
| 1* | 8.36 | 2.18 | 0.34 | 0.016 |
| 2 | 8.56 | 2.09 | 0.36 | 0.013 |
| 3 | 8.76 | 1.99 | 0.35 | 0.014 |
| 4 | 8.92 | 1.99 | 0.33 | 0.021 |
| 5 | 9.14 | 2.19 | 0.33 | 0.023 |
| 6 | 9.36 | 2.37 | 0.36 | 0.013 |
| 7 | 9.58 | 2.55 | 0.40 | 0.005 |
| 8 | 9.81 | 2.71 | 0.44 | 0.002 |
| 9 | 10.03 | 2.87 | 0.43 | 0.002 |
| 10 | 10.25 | 3.02 | 0.44 | 0.002 |
| 11 | 10.49 | 3.19 | 0.44 | 0.002 |

Note: Partial correlations controlling for infant age at scan, nonverbal IQ and parental education were reported. *This subsample included infants whose resting-state images were collected within the first year of life.

**Supplementary Table 9.** Partial correlation results between infant FC of the IFG module and word reading skills in subsamples ($n$ = 31) with increasing infant ages.

| **Subsamples** | **Mean of infant age (months)** | **SD of infant age (months)** | **Partial correlation *r* values** | ***p* values** |
|---|---|---|---|---|
| 1* | 8.79 | 1.67 | 0.40 | 0.036 |
| 2 | 9.04 | 1.68 | 0.42 | 0.025 |
| 3 | 9.26 | 1.72 | 0.41 | 0.030 |
| 4 | 9.47 | 1.76 | 0.41 | 0.029 |
| 5 | 9.79 | 2.08 | 0.43 | 0.022 |
| 6 | 10.11 | 2.33 | 0.45 | 0.017 |
| 7 | 10.45 | 2.56 | 0.46 | 0.014 |
| 8 | 10.77 | 2.77 | 0.46 | 0.014 |
| 9 | 11.10 | 2.96 | 0.43 | 0.021 |
| 10 | 11.41 | 3.13 | 0.39 | 0.039 |
| 11 | 11.74 | 3.33 | 0.40 | 0.034 |

Note: Partial correlations controlling for infant age at scan, nonverbal IQ and parental education were reported. *This subsample included infants whose resting-state images were collected within the first year of life.

**Supplementary Table 10.** Partial correlation results of FC between each of the four phonology-related ROIs and every module within the original language network in infancy with school-age behavioral outcomes, when controlling for infant age, nonverbal IQ, and parental education.

| | Oral language | Phonological skills | Word reading |
|---|---|---|---|
| FC between LIPL and IFG module | 0.10 | -0.03 | -0.10 |
| FC between LSMG and IFG module | -0.01 | -0.06 | -0.08 |
| FC between LSTG and IFG module | 0.13 | 0.10 | -0.09 |
| FC between RSTG and IFG module | 0.23 | 0.12 | -0.04 |
| FC between LIPL and MFG module | 0.01 | -0.05 | -0.33 |
| FC between LSMG and MFG module | 0.07 | 0.12 | -0.002 |
| FC between LSTG and MFG module | 0.02 | 0.19 | 0.13 |
| FC between RSTG and MFG module | 0.17 | 0.17 | 0.01 |
| FC between LIPL and TPG module | -0.04 | -0.06 | -0.03 |
| FC between LSMG and TPG module | 0.10 | 0.03 | 0.004 |
| FC between LSTG and TPG module | 0.12 | -0.14 | -0.04 |
| FC between RSTG and TPG module | -0.04 | -0.06 | 0.07 |

Note. Correlation coefficients were Pearson partial correlations between each pair of FC values and school-age behavioral outcomes controlling for infant age, nonverbal IQ and parental education. No significant associations were observed (all $p_{corrected} > 0.1$). LIPL: left inferior parietal lobule; LSMG: left supramarginal gyrus; LSTG: left superior temporal gyrus; RSTG: right superior temporal gyrus; IFG module: inferior frontal module; MFG module: middle frontal module; TPG: temporal and parietal module.

**Supplementary Table 11.** Hierarchical clustering and modularity analyses results based on sixteen ROIs including the original twelve language network ROIs and the four phonology-related brain regions.

| **Clustering results** | | **Clustering assignments of sixteen seed regions** | | | | | | | | | | | | | | | | **Modularity** |
|---|---|---|---|---|---|---|---|---|---|---|---|---|---|---|---|---|---|---|
| **Path length** | **N of Clusters** | **L IFG orb** | **L IFG** | **R IFG orb** | **R IFG** | **L MFG** | **R MFG** | **L Ant Temp** | **L Post Temp** | **R Ant Temp** | **R Post Temp** | **L AngG** | **R AngG** | **L IPL** | **L SMG** | **L STG** | **R STG** | **Q values** |
| 0.26 | 16 | 1 | 2 | 3 | 4 | 5 | 6 | 7 | 8 | 9 | 10 | 11 | 12 | 13 | 14 | 15 | 16 | 0.13 |
| 0.28 | 15 | 1 | 2 | 3 | 4 | 5 | 6 | 7 | 8 | 9 | 8 | 10 | 11 | 12 | 13 | 14 | 15 | 0.14 |
| 0.34 | 14 | 1 | 2 | 3 | 4 | 5 | 6 | 7 | 8 | 7 | 8 | 9 | 10 | 11 | 12 | 13 | 14 | 0.15 |
| 0.37 | 13 | 1 | 2 | 3 | 4 | 5 | 6 | 7 | 8 | 7 | 8 | 9 | 10 | 11 | 12 | 13 | 13 | 0.16 |
| 0.39 | 12 | 1 | 2 | 3 | 2 | 4 | 5 | 6 | 7 | 6 | 7 | 8 | 9 | 10 | 11 | 12 | 12 | 0.16 |
| 0.39 | 11 | 1 | 2 | 3 | 2 | 4 | 5 | 6 | 7 | 6 | 7 | 8 | 8 | 9 | 10 | 11 | 11 | 0.17 |
| 0.41 | 10 | 1 | 2 | 3 | 2 | 4 | 4 | 5 | 6 | 5 | 6 | 7 | 7 | 8 | 9 | 10 | 10 | 0.18 |
| 0.41 | 9 | 1 | 2 | 3 | 2 | 4 | 4 | 5 | 5 | 5 | 5 | 6 | 6 | 7 | 8 | 9 | 9 | 0.20 |
| 0.46 | 8 | 1 | 2 | 1 | 2 | 3 | 3 | 4 | 4 | 4 | 4 | 5 | 5 | 6 | 7 | 8 | 8 | 0.21 |
| 0.48 | 7 | 1 | 2 | 1 | 2 | 3 | 3 | 4 | 4 | 4 | 4 | 5 | 5 | 6 | 6 | 7 | 7 | 0.21 |
| 0.50 | 6 | 1 | 2 | 1 | 2 | 3 | 3 | 4 | 4 | 4 | 4 | 5 | 5 | 6 | 6 | 4 | 4 | 0.25 |
| 0.69 | 5 | 1 | 1 | 1 | 1 | 2 | 2 | 3 | 3 | 3 | 3 | 4 | 4 | 5 | 5 | 3 | 3 | 0.27 |
| 0.73 | 4 | 1 | 1 | 1 | 1 | 2 | 2 | 3 | 3 | 3 | 3 | 3 | 3 | 4 | 4 | 3 | 3 | 0.26 |
| 0.77 | 3 | 1 | 1 | 1 | 1 | 2 | 2 | 3 | 3 | 3 | 3 | 3 | 3 | 1 | 1 | 3 | 3 | 0.25 |
| 0.88 | 2 | 1 | 1 | 1 | 1 | 2 | 2 | 1 | 1 | 1 | 1 | 1 | 1 | 1 | 1 | 1 | 1 | 0.14 |

Note: Red fonts indicate the modular partition with the maximum modularity Q value.

**Supplementary Table 12.** Partial correlation results between infant FC of each module within the extended language network and behavioral outcomes controlling for infant age, nonverbal IQ, and parental education.

| **Module** | **IFG** | **MFG** | **Temporal** | **AngG** | **Parietal** |
|---|---|---|---|---|---|
| **Phonological skills** | 0.38* | -0.13 | -0.08 | 0.07 | -0.002 |
| **Oral language** | 0.11 | 0.01 | 0.02 | 0.01 | -0.03 |
| **Emergent word reading** | 0.43* | 0.03 | 0.04 | 0.15 | 0.05 |

Note: IFG module: inferior frontal module; MFG module: middle frontal module; AngG module: angular gyri module.

*$p_{corrected} < 0.05$

**Supplementary Table 13.** Two-sample *t*-test results between two cohorts of infants with different imaging acquisition sequences

| Characteristics at different stages | Group | *n* | Mean | SD | *t*-value | df | *p*-value |
|---|---|---|---|---|---|---|---|
| ***Infancy Imaging Time Point*** | | | | | | | |
| **Scanning age (months)** | 1 | 43 | 10.3 | 3.7 | 3.12 | 74 | 0.003 |
| | 2 | 33 | 7.9 | 2.9 | | | |
| **MSEL receptive language** | 1 | 28 | 44.4 | 8.4 | -0.77 | 59 | 0.443 |
| | 2 | 33 | 46.1 | 8.7 | | | |
| **MSEL expressive language** | 1 | 35 | 46.5 | 7.6 | -2.62 | 66 | 0.011 |
| | 2 | 33 | 51.9 | 9.5 | | | |
| ***Kindergarten-age Language skills*** | | | | | | | |
| **Test age (months)** | 1 | 43 | 75.4 | 10.9 | 6.61 | 74 | $< 0.001$ |
| | 2 | 33 | 61.8 | 4.9 | | | |
| **Phonological skills** | 1 | 38 | 106.1 | 15.8 | 1.10 | 59 | 0.274 |
| | 2 | 23 | 101.4 | 16.6 | | | |
| **Oral language** | 1 | 43 | 114.5 | 12.9 | 0.80 | 74 | 0.424 |
| | 2 | 33 | 111.9 | 15.1 | | | |
| ***Emergent Reading Abilities*** | | | | | | | |
| **Test age (months)** | 1 | 36 | 100.5 | 11.3 | 5.14 | 39 | $< 0.001$ |
| | 2 | 5 | 74.3 | 2.5 | | | |
| **Test of Word Reading Efficiency** | 1 | 36 | 104.9 | 16.2 | 0.81 | 39 | 0.422 |
| | 2 | 5 | 98.7 | 12.8 | | | |

Group 1: infants' resting state images were acquired before 2017 using a single-band sequence with a TR of 3s;
Group 2: infants' resting state images were acquired in 2017 and after using a simultaneous multi-slice (SMS) sequence with a TR of 0.95s.

**Supplementary Table 14.** Partial correlation results between FC of each module within the infant language network and behavioral outcomes controlling for infant age at scan, nonverbal IQ, parental education, and testing ages of the follow-up behavioral assessments.

| | **IFG module** | **MFG module** | **TPG module** |
|---|---|---|---|
| **Phonological skills** | $r = 0.38^*$, $p = 0.004$ | $r = -0.13$, $p = 0.352$ | $r = -0.06$, $p = 0.657$ |
| **Oral language** | $r = 0.11$, $p = 0.360$ | $r = -0.01$, $p = 0.916$ | $r = -0.14$, $p = 0.234$ |
| **Emergent word reading** | $r = 0.45^*$, $p = 0.006$ | $r = 0.03$, $p = 0.846$ | $r = 0.12$, $p = 0.497$ |

Note: IFG module: inferior frontal module; MFG module: middle frontal module; TPG module: temporal and parietal module. $^*p_{corrected} < 0.05$

**Supplementary Table 15.** Partial correlation results between infant FC of the IFG module and subsequent phonological skills in subsamples (*n* = 47) with increasing ages of behavioral assessments of phonological skills.

| **Subsamples** | **Mean of age (years)** | **SD of age (years)** | **Partial correlation *r* values** | ***p* values** |
|---|---|---|---|---|
| 1* | 5.55 | 0.63 | 0.42 | 0.005 |
| 2 | 5.61 | 0.65 | 0.42 | 0.005 |
| 3 | 5.66 | 0.67 | 0.42 | 0.005 |
| 4 | 5.71 | 0.69 | 0.38 | 0.010 |
| 5 | 5.75 | 0.70 | 0.37 | 0.013 |
| 6 | 5.80 | 0.71 | 0.43 | 0.004 |
| 7 | 5.85 | 0.73 | 0.42 | 0.004 |
| 8 | 5.90 | 0.74 | 0.41 | 0.006 |
| 9 | 5.95 | 0.75 | 0.40 | 0.007 |
| 10 | 6.00 | 0.77 | 0.37 | 0.013 |
| 11 | 6.06 | 0.78 | 0.35 | 0.020 |
| 12 | 6.11 | 0.79 | 0.31 | 0.038 |
| 13 | 6.17 | 0.80 | 0.32 | 0.034 |
| 14 | 6.22 | 0.82 | 0.34 | 0.026 |

Note: Partial correlations controlling for infant age at scan, nonverbal IQ and parental education were reported. *This subsample included participants whose phonological skills were assessed before 7 years old.

**Supplementary Table 16.** Partial correlation results between infant FC of the IFG module and emergent word reading skills in subsamples ($n$ = 33) with increasing ages of behavioral assessments of reading skills.

| **Subsamples** | **Mean of age (years)** | **SD of age (years)** | **Partial correlation *r* values** | ***p* values** |
|---|---|---|---|---|
| 1* | 7.76 | 0.96 | 0.50 | 0.005 |
| 2 | 7.85 | 0.93 | 0.52 | 0.003 |
| 3 | 7.94 | 0.90 | 0.54 | 0.002 |
| 4 | 8.03 | 0.85 | 0.51 | 0.004 |
| 5 | 8.13 | 0.81 | 0.50 | 0.004 |
| 6 | 8.21 | 0.78 | 0.50 | 0.005 |
| 7 | 8.32 | 0.80 | 0.50 | 0.005 |
| 8 | 8.43 | 0.81 | 0.52 | 0.003 |

Note: Partial correlations controlling for infant age at scan, nonverbal IQ and parental education were reported. *This subsample included participants whose word reading skills were assessed before 9 years old.
